# Global evolutionary patterns of *Yersinia pestis* and its spread into Africa

**DOI:** 10.1101/2024.11.26.625443

**Authors:** Guillem Mas Fiol, Frédéric Lemoine, Damien Mornico, Guillaume Bouvier, Aida Andrades Valtueña, Sebastian Duchene, Pascal Campagne, Charlotte Balière, Aurélia Kwasiborski, Valérie Caro, Rémi Beau, Cyril Savin, Manuel Céspedes, Minoarisoa Rajerison, Jean-Christophe Shako, Elisabeth Carniel, Philip Slavin, Nicolás Rascovan, Javier Pizarro-Cerdá

## Abstract

The zoonotic pathogen *Yersinia pestis*, the etiologic agent of plague, has caused three major pandemics and diversified in different lineages currently established in endemic areas worldwide^1–3^. However, some regions like continental Africa have been poorly covered within the global diversity and epidemiological history of this pathogen^2,4–6^. Here, we report the whole-genome sequences of 1,124 *Y. pestis* isolates collected from endemic areas worldwide over 116 years, nearly doubling the available genomic data for the species. By integrating population genomics and historical research, we retrace the introduction of multiple *Y. pestis* lineages into continental Africa, revealing the diversity of the 1.ANT lineage, its historical emergence and its spread to and within Africa since the late 17th century. We identify key mechanisms of genome evolution, including signatures of adaptive evolution present in virulence and biofilm-related genes such as RovA, a master virulence regulator, which likely play a role in the pathogen’s adaptation and endemic persistence. Additionally, our findings reveal an increased trajectory of genome degradation and expansion of IS elements in different lineages. This trend appears especially pronounced in 1.ANT genomes, promoting the remarkable genomic variation within this lineage. Taken together, our findings shed light on the introduction and evolutionary history of plague in Africa and provide a comprehensive framework for understanding the global diversity and genome evolution of *Y. pestis*, revealing potential factors contributing to its long-term adaptation in endemic areas.

---

Plague is a fatal zoonotic disease caused by the gram-negative bacterium *Yersinia pestis*. The pathogen is transmitted to humans mainly by fleas from rodents, which are its primary reservoir^1^. *Y. pestis* has infected humans for at least 5,000 years^7,8^, claiming millions of human lives across three major pandemics that have profoundly impacted human history^2^. Reemerging and spreading out of Asia on multiple occasions, plague is maintained in areas known as endemic foci, currently present in Africa, Asia, the easternmost extremes of Europe (i.e.: Caucasus), and the Americas^9^. Recently, *Y. pestis* has also been classified as a priority pathogen for pandemic research preparedness by the WHO^10^.

Considered as one of the most virulent bacterial pathogens to humans, *Y. pestis* is a recently emerged clone of its enteropathogenic ancestor *Y. pseudotuberculosis*^11^ that evolved through a series of gene gain and genome reduction events^12,13^. These changes allowed *Y. pestis* to adapt to an insect vector and to establish new transmission routes to infect mammal hosts^14^. In the recent years, the steady expansion of genomic data has allowed to retrace the evolution of *Y. pestis*, leading to the identification of 20 major modern lineages from the sequencing of bacterial isolates circulating in the past decades^4,15^. Furthermore, unprecedented insights into the phylogeography of extinct lineages that spread during historical plague pandemics and outbreaks up to millennia ago have been uncovered from the analysis of ancient DNA samples^7,8,16^. However, the overall portrayal of the global genetic diversity and evolution of *Y. pestis* is biased towards genomes from isolates recovered mainly from areas across Eurasia^4,5,15^. Extensive genomic surveillance has been conducted in only a handful of plague endemic areas, such as Madagascar^17^ and China^18,19^, while other active regions remain underrepresented. This is illustrated by the scarcity of genomes from continental (mainland) Africa, despite the highest number of human plague cases in the last decade have been reported from this continent after recurrent episodic outbreaks in the Democratic Republic of Congo (DRC)^20^. For instance, as few as four genomes are available for the 1.ANT lineage, which is believed to be restricted to Central-Eastern Africa^4,21^, preventing the implementation of phylogenetic and temporal frameworks to retrace the introduction and evolution of this lineage into Africa^22^. Moreover, previous studies on the intraspecific evolution of *Y. pestis* have essentially focused on mutation rates within core genes^5,6^, while other genomic mechanisms driving strain-to-strain genome variation and their role in the adaptation of this pathogen in endemic regions remain poorly explored^19,23^.

Here, we take steps to overcome these limitations by sequencing the genomes of more than a thousand isolates from epidemiologically relevant endemic regions worldwide, including Africa, that were collected from the turn of the 20^th^ through the early 21^st^ centuries. We use population genomics methods to develop the most extensive analysis to date of *Y. pestis* diversity and we take an integrative approach to infer the emergence and introduction of lineages that have caused plague in continental Africa in the past centuries. Finally, we combine different species-wide genomic analyses to identify the main genetic mechanisms driving the functional diversification and potential adaptation of this pathogen across endemic areas.

## Results

### Sequencing of a worldwide *Y. pestis* collection

To strengthen our understanding of the genetic diversity of contemporary *Y. pestis*, we selected 1,124 *Y. pestis* isolates from the historical collection of the Institut Pasteur in Paris, which were collected from 29 different countries covering important plague endemic areas in the world, including Africa, the Middle East, South-East Asia and South America (Supplementary Fig. 1 and Supplementary Table 1). Notably, the collection includes 161 *Y. pestis* isolates from continental Africa, while the genomes of only 6 isolates were previously available in public databases (Supplementary Table 2). The collection comprises bacterial samples isolated over 116 years (1899-2013), with the oldest sample closely following the first ever isolation of *Y. pestis* in 1894. Isolates had been collected mostly from human clinical cases of plague (n=793, 70.5%). The collection also includes isolates from animal plague surveillance efforts in endemic countries, including a range of rodent species (n=164, 14.6%) and fleas (n=37, 3.3%) (Supplementary Fig. 1, Supplementary Table 1). DNA was extracted from the live culture of each isolate and was shotgun sequenced to generate high genome coverage and *de novo* assemblies. Of note, no antimicrobial resistance determinants were detected when querying the 1,124 assemblies against ResFinder 4.0 database.

### The updated global diversity of *Y. pestis*

To place the new genomes in relation to the global diversity of the species, we conducted a phylogenetic analysis including most of the publicly available modern and ancient genomes of *Y. pestis* (1,682 genomes) together with the 1,124 novel sequences (Supplementary Table 2). We identified core genome variants (28k variants over 3.62 million positions) and used them to generate a maximum likelihood phylogeny including a total of 2,806 genomes, building up the largest and most comprehensive species tree of *Y. pestis* to date (**Fig.1a,** Extended Data Fig. 1). The tree classifies the genomes into the previously described 20 main lineages of uneven frequency and geographic distribution (Supplementary Fig. 2), named according to the established nomenclatures^4,5,15^. As expected from the geographic distribution of strains, most of the new genomes (90%) clustered within the 1.ORI lineage, which displays a star-like topology reflecting its recent expansion during the so-called third plague pandemic^4^. This branch appears particularly populated after different episodes of extended sampling in various endemic countries such as Madagascar, Vietnam and Brazil across the past decades (**Fig. 1bc**). The enrichment of 1.ORI genomes allowed us to identify two new monophyletic clades: 1.ORI4 in South-East Asia (Vietnam, China, Myanmar); and 1.ORI5, which spread across South America, Europe and Africa, reaching, for the latter, Morocco, Senegal, Algeria, South Africa, Namibia and Zimbabwe (Extended Data Fig. 2).

**Figure 1.**
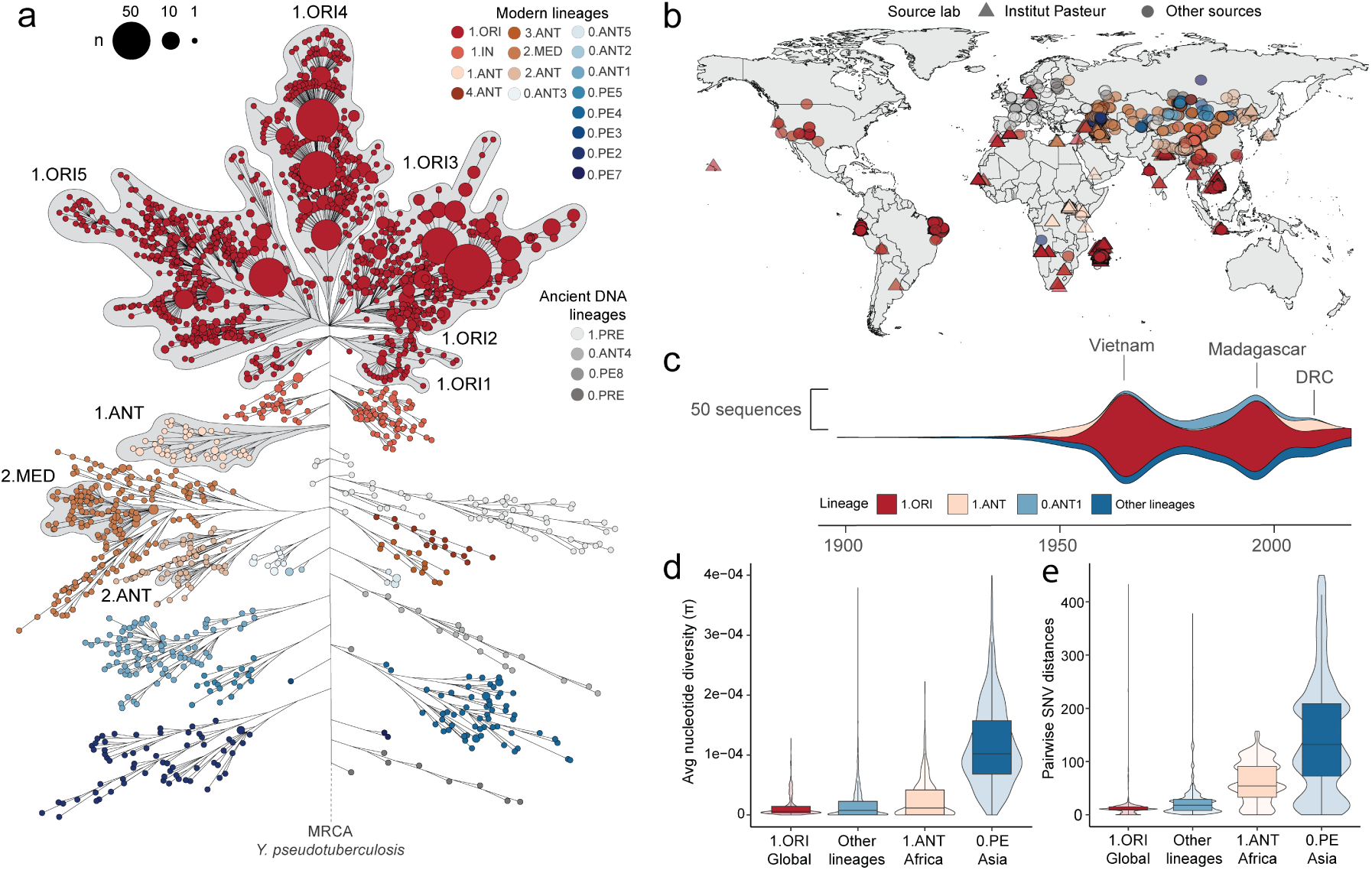
Global genetic diversity of *Y. pestis* species. **a**, Maximum likelihood phylogeny of 2,806 Y*. pestis* modern and ancient genomes. Shaded areas surrounding branches and labels indicate branches including new sequences from the Institut Pasteur collection (a fully annotated tree is displayed in Extended Data Fig.1). The tree is scaled on the number of substitutions per genomic site. Branches with lengths shorter than 1e-7 were collapsed and lengths were log-scaled to improve visualization of lineages. **b**, Geographic distribution of sequences included in the phylogeny, colored according to the same lineages in panel a. **c**, Steamgraph showing the temporal changes in *Y. pestis* lineages sampling within the dataset of modern genomes. Labels mark periods of increased sampling associated with the epidemiological context of three endemic countries. The complete metadata of new sequences is shown in Supplementary Table 1. **d**, Average nucleotide diversity (*π*) within modern lineages (n=2,722 genomes). Estimates were computed by 10-kb windows from the variant tables (VCF). **e**, Distribution of pairwise SNV distances between genomes of modern lineages.

Our dataset also significantly increases the sampling of two other *Y. pestis* lineages: 2.MED and 1.ANT (Supplementary Fig. 3). Three isolates from an outbreak in Northern Africa (Libya)^24^ fall within a 2.MED1 clade that includes new genomes from the Middle East (Iran, Turkey) and the Caucasus (Russia, Azerbaijan). For 1.ANT, 42 new continental African sequences and an isolate from Yemen populate the three different clades of this lineage (1.ANT1, 1.ANT2 and 1.ANT3)^4^, which was previously represented by just four genomes from Central-Eastern Africa (**Fig. 1ab**). The lineage is characterized by a higher sampling frequency in the recent decades, associated with plague outbreaks in the DRC and Uganda^21,25^ (**Fig. 1c**). Our new dataset thus recapitulates the worldwide diversity of *Y. pestis* species and fills a former gap in genomes from continental Africa, opening the opportunity to explore the evolutionary and spreading trajectories of the pathogen across this region.

### High genetic diversity of *Y. pestis* in Africa

To better understand the diversification of contemporary *Y. pestis* populations, we estimated the nucleotide diversity (*π*) within modern *Y. pestis* genomes, rendering varied levels of genetic diversity between lineages (Kruskal-Wallis test, *p*<0.001, **Fig. 1d**). Consistent with their ancestral position and early divergence in the *Y. pestis* phylogeny, 0.PE (Pestoides) genomes from Asia have accumulated the highest diversity. By contrast, the globally distributed 1.ORI lineage has accumulated little diversity given its recent emergence and clonal expansion in the context of the third plague pandemic^4,5^. Intriguingly, African 1.ANT genomes display some of the highest average *π* values despite their relatively recent emergence as part of branch 1 (**Fig. 1d**). Complementing this observation, the distances in single nucleotide variants (SNV) separating pairs of isolates within each lineage are also within the highest in 1.ANT lineage (Wilcoxon-Mann-Whitney test, *p*<0.05, **Fig. 1e**, Supplementary Table 3), suggesting that a substantial fraction of its diversity has accumulated since its early diversification in Africa. Overall, our results highlight a high genetic diversity of *Y. pestis* in continental Africa, as evidenced by the presence of three independent lineages (1.ORI, 2.MED and 1.ANT) and the remarkable diversity accumulated within the 1.ANT lineage since its arrival to Central-Eastern Africa.

### History of *Y. pestis* spread in continental Africa

To provide a temporal framework of plague introductions into continental Africa, we have analyzed both new and previously published genomes and reconstructed a time-calibrated phylogeny using BEAST2 (Supplementary Fig. 4, Supplementary Table 4). We first investigated the arrival of the 1.ORI5 clade into Africa, where it subsequently diverged into two subclades, one associated with Western Africa and the other with Southern Africa (Extended Data Fig. 3). The inferred dates for the diversification of both subclades were estimated around 1900 (95% highest posterior density (HPD) intervals: 1802-1894, median 1852; and 1843-1934, median 1891, respectively), at the onset of the third plague pandemic, when plague spread through maritime routes^4^ (**Fig. 2ab**, Supplementary Table 5). This is corroborated by historical records, indicating the arrival of the plague in South-East African ports in 1898-9, before spreading further inland (Namibia in 1931 and Zimbabwe in 1972^25,26^, see Supplementary Information 1). Conversely, the West African 1.ORI5 subclade may have emerged even prior to its initial introduction and focalization in Maghreb (1899-1909), as inferred from our phylogenetic analysis^27,28^. Furthermore, we estimated the introduction of the 2.MED lineage into Northern Africa in the early 20^th^ century, as inferred by the date of the most recent common ancestor (MRCA) of the Libyan genomes (95% HPD: 1911-1947, median 1937; **Fig. 2a**). The three Libyan isolates were collected during the 2009 outbreak and fall within a larger 2.MED1 clade that likely emerged a century earlier (95% HPD: 1749-1865; median date: 1812). This 2.MED1 clade appears to have emerged and focalized in Kurdistan (or possibly in Iranian Azerbaijan), subsequently radiating into neighboring Caucasus, Anatolia and Iran, as indicated by the phylogenetic analysis (**Fig. 2b**, Extended Data Fig. 3), but likely into Iraq too, as the historical records suggest (Supplementary Information 1).

**Figure 2.**
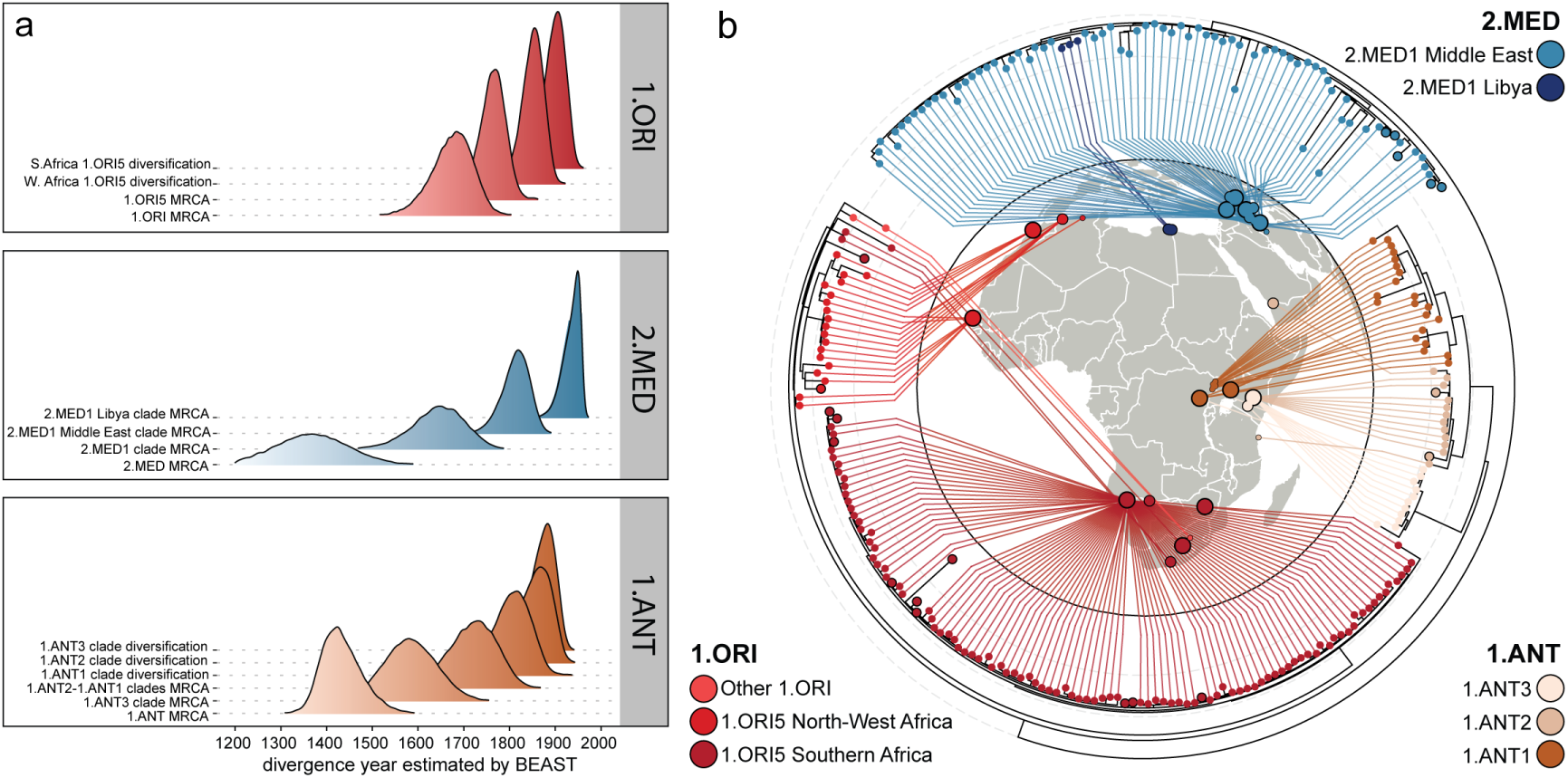
Phylogeography of *Y. pestis* in Africa. **a**, Distribution of posterior divergence dates for nodes in the time-calibrated phylogenetic tree inferred using BEAST2. MRCA nodes represent the last node a monophyletic clade shares with other clades, while diversification nodes represent the node after which extant isolates diversify. For the complete maximum clade credibility tree including all lineages, see Supplementary Fig. 4. **b**, Maximum-likelihood phylogeny of genomes from three distinct lineages introduced into continental Africa. The tree was extracted from the complete species tree shown in Fig. 1. Each genome in the tree is linked to the available geographic location in Africa, with the smallest circles indicating town-level precision, intermediate size indicating county-level information and largest circle sizes when only information of the country of isolation is available.

The deepest branching lineage in Africa is 1.ANT, diverged off Branch 1B, which has been associated with the *pestis secunda* in West Eurasia and North Africa (1356-66)^6,16,29^. Previous works aiming to date the emergence of 1.ANT have been so far hampered by the scarcity of available genomes, leading to irreproducible estimates^22^. In contrast with previous studies^5,22^, our molecular clock analyses estimate narrower time frames for the MRCA of 1.ANT, which emerged in medieval times (95% HPD 1355-1528; median date: 1429, **Fig. 2a**, Extended Data Fig. 3). We also inferred the MRCA dates of the different 1.ANT clades, revealing that they all diverged between the 16^th^ and 18^th^ Centuries (1.ANT3: 95% HPD 1472-1697; median date: 1582; 1.ANT2-1.ANT: 95%HPD 1603-1810; median date: 1714; **Fig. 2a**, Extended Data Fig. 3, Supplementary Table 5). To elucidate the chronology of its introduction and prevalence in East Africa, we conducted a thorough investigation of historical records (Supplementary Information 1). As all currently described lineages ancestral to 1.ANT have a Eurasian origin, we hypothesized the emergence of 1.ANT to have occurred outside of East Africa. We explored all potential routes of introduction into Africa and found records supporting the arrival into the Great Lakes region of East Africa, most likely from Egypt via the Nile, around 1700. This introduction was likely favored by famine- and conflict-induced migration linked to a short-term climatic anomaly reported during that period (Supplementary Information 1). We then identified the earliest historical records reporting a disease matching plague in the Kingdom of Buganda (northwestern shore of Lake Victoria), which were followed by successive reports of plague outbreaks limited to this region for over a century. The phylogenetic tree of 1.ANT indicates an early diversification with the emergence of 1.ANT3, which has then evolved locally for at least two centuries without further branching events in its phylogeny (Extended Data Fig. 3). Thus, both the historical records and the phylogenetic analyses are consistent with an initial diversification of the 1.ANT restricted to the Buganda territory.

From 1840 or so, however, the historical records indicate a radiation of plague into other areas around the Great Lakes, eventually reaching the Swahili Coast of the Indian Ocean by the end of the 19^th^ century. Complementarily, our phylogenetic analysis reveals that the extant strains within each clade of 1.ANT diversified at this exact same period (**Fig. 2a**). This was a period characterized by increasing commercialization of the region and its integration into the Indian Ocean system, which was facilitated first by Arab merchants, and later by European settlers, colonial authorities, traders and missionaries. In addition, this period witnessed an increased intensity in conflicts between local polities, as well as inter-regional migration around the Great Lakes. Our genomic and historic findings are therefore consistent with an expansion of plague across East Africa, within the shifting context of the 19^th^ century in the region, reaching Uganda, East Congo and Tanzania, and spreading further into South Arabia, as evidenced by the isolate from Yemen falling within the 1.ANT2 clade (**Fig. 2ab**, Supplementary Information 1). The integrated analysis of genomic and historical evidence demonstrates at least three centuries of plague presence in Africa, with 1.ANT representing the first lineage to establish a long-term, endemic reservoir outside of Eurasia.

### Detecting adaptive signals in *Y. pestis*

*Y. pestis* evolution is thus characterized by multiple introduction events into different geographic areas followed by its local diversification^4,5^, where the pathogen circulates under contrasting environments and animal reservoirs^2^. The substantial increase in the number of genomes, combined with the diverse geographic sources of the compiled dataset, offers new opportunities to detect hallmarks of adaptation and functional diversification within *Y. pestis* genomes. To explore this question, we first looked at genes presenting an excess of non-synonymous (nonSyn) or homoplastic mutations (emerging independently multiple times), two features associated with potential positive selection^5,30^. By evaluating 15,962 single nucleotide variants (SNVs) called in modern genomes (Methods), we detected 60 genes with a number of nonSyn mutations significantly deviating from theoretical expectations under neutral evolution (**Fig. 3a**, Supplementary Table 6). Over a third of these genes (36%) also present convergent (homoplastic) non-Syn mutations that independently emerged at least three times across the phylogeny (**Fig. 3b**, Supplementary Table 7), suggesting adaptive evolution may be driving their functional diversification. Among the 60 loci, only 7 (*aspA*, *rpoZ*, *sspA*, *purR*, *gcvA, YPO0623* and *YPO2210*) had been previously reported for their increased rates of nonSyn mutations and/or homoplasies in *Y. pestis*^5,19,31^. The predicted effect of the mutations also revealed a large proportion of nonsense mutations within some of the loci (*cytR*, *rnd*, *YPO0623*, *pldA*, *rpoS* and *YPO2444*), suggesting that beyond amino acid changes, adaptive pressures may be driving their loss of function (Supplementary Fig. 5). Notably, the SNVs found in the 60 identified genes appear overall scattered throughout the phylogeny, while few (13%) have been fixed in complete clades (Extended Data Fig. 4). Furthermore, upon closer examination of the geographic distribution of isolates carrying mutations in each of the 60 loci, we identified two genes for which most (>90%) of the mutated isolates had an African origin, indicating the emergence of these variants occurred mainly in Africa (Supplementary Fig. 6). These genes are *purR*, coding a direct regulator of the expression of virulence genes in pathogenic *Staphylocccus aureus*^32^; and *rpsC*, coding a ribosomal subunit protein.

**Figure 3.**
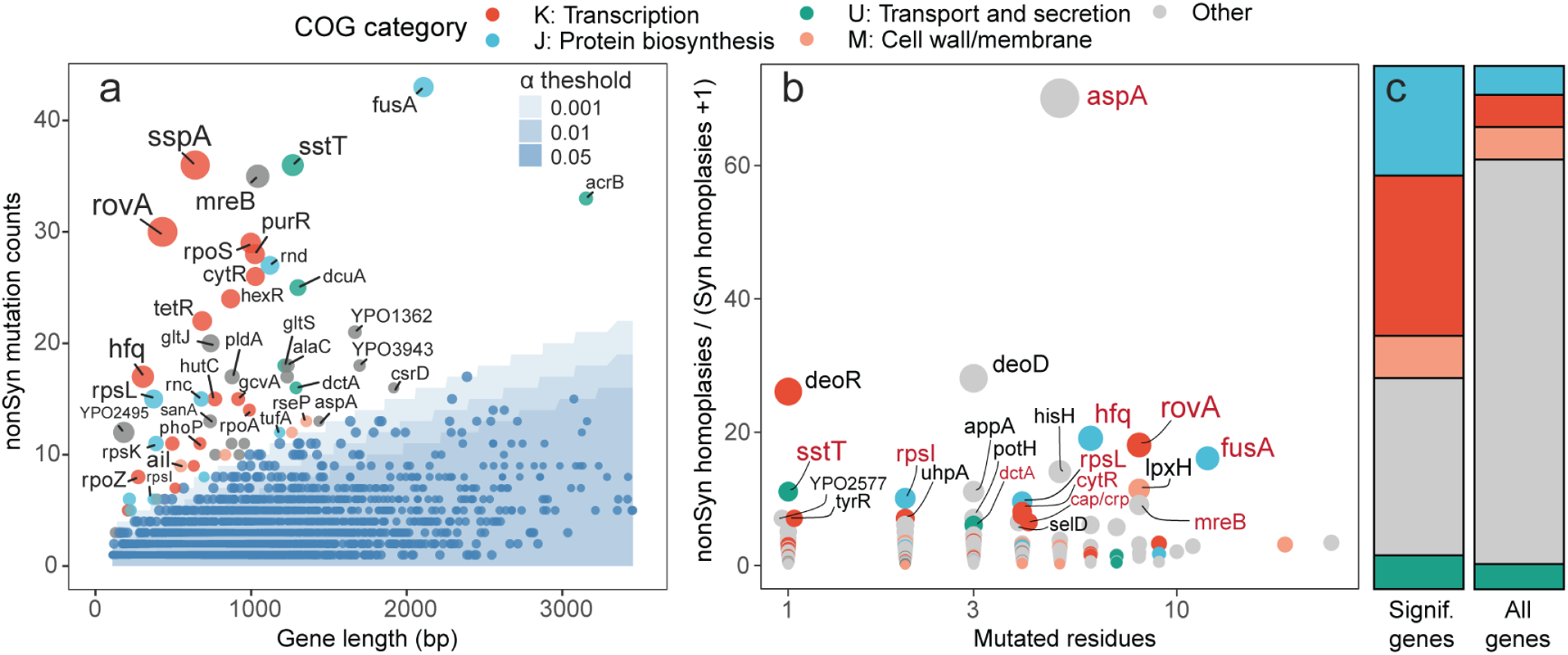
Genes with adaptive signatures in *Y. pestis* evolution. **a**. Identification of genes (n=60) with significantly higher mutation densities using a sequential Bonferroni-corrected *χ*^2^ goodness of fit test. Each circle represents a gene, for which the total number of nonSyn mutations detected is displayed as function of its sequence length. Dots are colored according to functional categories defined by Clusters of Orthologous Genes (COG). Circle sizes are proportional to the contribution of each gene to the deviation from the expected number of nonSyn mutations (contribution to the *χ*^2^ goodness-of-fit statistic). The expected number of mutations at each gene length based on different *α* levels (0.05, 0.01, 0.001) are indicated in shades of blue. For the complete list of genes diversifying at higher rates and their annotations, see Supplementary Table 6. **b**, Genes with highest ratio of nonSyn mutation events emerging independently through the phylogeny (homoplasies). Scatterplot displaying the rate of homoplastic nonSyn mutations per gene normalized to the total number of syn mutation events per gene, against the total number of nucleotide sites affected by homoplasies. Each dot represents a gene, and dots are colored according to the same functional categories as in panel a. Top-ranking genes regarding their ratio of nonSyn homoplastic mutation events are labelled, and red labels indicate highly variable genes also identified in panel a. For the detailed ratios of homoplastic mutation events within the complete gene set, refer to Supplementary Table 7. **c**, Enrichment of Transcription (K) and Translation (J) functional categories in genes diversifying at higher rates compared to the total set of genes (Fisher’s exact test, *p*<0.001 for both categories).

To gain insights into the functional roles of the identified genes diversifying at higher rates, we classified all protein-coding genes according to the different functional categories defined by the Clusters of Orthologous Groups (COGs). Interestingly, we observed a significant enrichment in functions related to transcription and protein biosynthesis (Fisher’s exact test, *p*<0.001) in the set of genes with higher mutation densities when compared to all other genes (**Fig. 3c**, Supplementary Table 8). The top-ranking genes identified include global gene expression regulators such as *rovA*, *phoP*, *sspA* and *hfq*, implicated in the control of wide-range functions including virulence, biofilm formation and stress response^33–36^. Several of the identified factors have also been associated (see Supplementary Table 6 for the associated references), in *Yersinia* spp. and other bacterial pathogens, with virulence (*ail, fusA*, *purR*, *cytR*, *aspA*), response to stress (*rpoB*, *rpoC*, *sstT*, *dctA*, *YPO0623*), biofilm formation (*rpoS*, *rpoZ*, *rpoA*) and antimicrobial resistance (*rpsL*). Since regulatory genes often influence a broad range of functions, the variants in some of the identified factors may be associated with responses to similar conditions or stresses. This is supported by the detection of positive correlations (*r*>0.5) between 19 of the highly variable genes in terms of variant co-occurrence across the analyzed genomes (Extended Data Fig. 5), suggesting that shared ecological conditions may be driving the coupled diversification of some of the genes. Taken together, our analysis reveals a signature of increased diversification in regulatory factors known to modulate relevant aspects of *Y. pestis* infection and transmission cycle.

### Adaptive mutations in RovA master virulence regulator

We then examined whether the variants found in genes with an excess of mutations were randomly distributed across their sequence (i.e., without an evident functional significance), or rather more frequently observed at amino acid residues that modulate protein functions. To test this hypothesis, we focused on *rovA/slyA* gene, which deviates the most from theoretical expectations of protein-altering variants per gene and has enough mutations in the genomic dataset for proper evaluation (**Fig. 3a**). This locus encodes a global transcriptional regulator of key factors involved in virulence and biofilm formation in pathogenic *Yersinia* spp.^33,37,38^. A total of 30 nonSyn mutations were found all along the length of its sequence, 9 being homoplastic and 6 nonsense (Supplementary Fig. 7, Supplementary Table 9). Most RovA variants show low frequency in *Y. pestis* lineages, with the exception of H22P substitution that is highly prevalent in 1.ANT2 and 1.ANT1 genomes, indicating that it emerged and became fixed in these African clades (**Fig. 4a**, Supplementary Table 9). Since RovA transcriptional activity requires binding to DNA promoter sequences as well as its own dimerization, we wondered whether the identified variants would likely impact such interactions. To explore this question, we mapped the mutated residues observed across the *Y. pestis* genomes on the resolved structure of RovA interacting with its DNA promoter fragment^39^ (**Fig. 4b**). Strikingly, we found an enrichment of mutations in residues which are at a close distance (≤ 4 Å) to the target DNA molecule, allowing for hydrogen bond interactions (**Fig. 4c**). Eleven nonSyn mutations indeed fall in known DNA-binding residues of RovA, including Q49, E59, V64, D68, R78 and R85^39,40^ (Supplementary Table 9). In addition, we found two mutations (E113*, S119*) that generate a premature stop in the C-terminal domain of RovA, in which deletions are known to impair its dimerization and transcriptional activation activity^40^. These findings overall provide compelling evidence that selective pressures have shaped the evolution of RovA by acting on functionally relevant sites.

**Figure 4.**
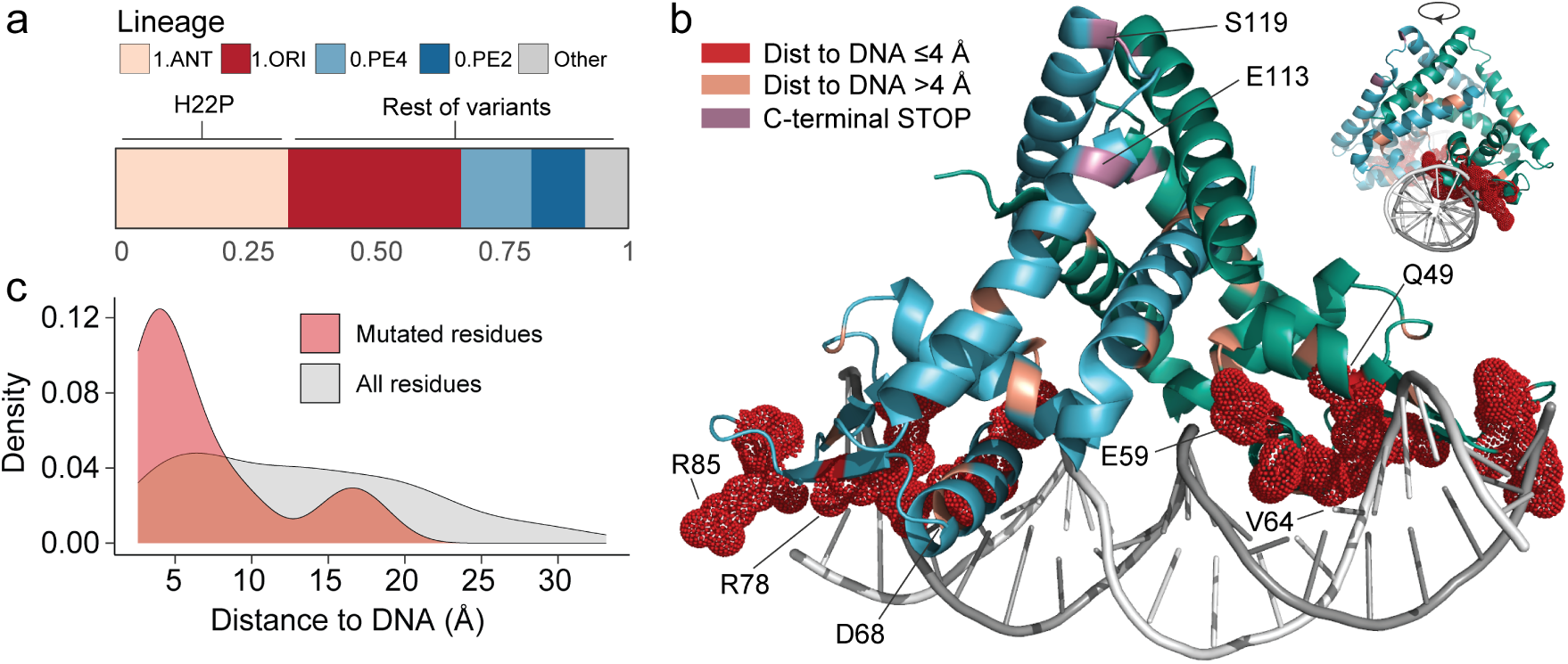
Enrichment of variants close to DNA in RovA global transcriptional regulator. **a**, Lineage distribution of genomes carrying one or more of the 30 nonSyn variants observed in *rovA* locus (see Supplementary Table 9 for details). H22P variant has been fixed in 1.ANT1-1.ANT2 genomes. **b**, Mapping of the identified mutated residues on the resolved structure of RovA dimers interacting with RovA promoter DNA molecule (PDB ID: 4AIJ). The surfaces of mutated residues which are at an estimated distance to DNA ≤ 4Å are shown as red dots. Labels indicate mutated residues known to be implicated in binding of RovA to target promoter DNA^39,40^ (Q49, E59, V64, D68, R78, R85), and C-terminal residues in which premature STOP codon mutations impact the dimerization region required for its activity (E113, S119). **c**, Closest distance to DNA molecule estimated for each residue in RovA structure, showing an enrichment of mutations in residues at close distance to DNA (4Å) in *Y. pestis* populations.

### Extensive gene loss and IS expansion in African strains

Besides adaptive mutations in conserved genes, the distinct evolutionary paths taken by *Y. pestis* lineages following their local evolution in different regions may have also led to variations in gene content. To address this question, we investigated the trajectories of gene gains and gene losses in *Y. pestis*. We first searched for likely acquired genes by querying more than 2,000 modern genomes against the MOB suite database of plasmids and found only 21 elements, including 9 novel plasmids, a putative chromosomal integrative and conjugative element (ICE) and a putative phage (Supplementary Tables 10 and 11). There were no unifying features on the likely acquired replicons, although 8 plasmids contain gene clusters associated with type IV secretion systems (Supplementary Fig. 8). Interestingly, the distribution of the elements along *Y. pestis* phylogeny revealed the parallel acquisition of a ∼22-k base (kb) cryptic plasmid (pCRY) in African isolates from 1.ANT2 and 1.ORI5 clades, a plasmid previously identified exclusively in a few Asian isolates^41^. The putatively acquired sequences are generally <50-kb in length, display a broad range of GC content and share homology with sequences of other Enterobacterales species, suggestive of their acquisition through horizontal gene transfer (Extended Data Fig. 6). However, the low prevalence of these elements, present only in 0.03-3.6% of total isolates, suggests that gene gain occurs infrequently in *Y. pestis*, likely playing a limited role in shaping gene content differences between lineages.

We thus considered another possibility, being that major gene content differences between lineages may rather be a consequence of gene loss events. A scan of chromosomal regions lacking mapping reads in the complete genomic dataset allowed us to generate a detailed cartography of large deletions *Y. pestis* evolution (**Fig. 5a**). We identified a total of 20 deleted regions, from which only 7 had been previously reported^6,12,42^ (Extended Data Table 1). The deletion islands (DI) have between 8,5 and 100-kb (median ∼30-kb) in length and in total sum up to 780-kb (17% of the reference chromosome), representing a substantial fraction of the genome that would be expendable in the species. The phylogenetic distribution of each DI revealed the parallel independent loss of several of them during *Y. pestis* evolution (Extended Data Fig. 7), suggesting their loss may be adaptive under certain conditions. One of such examples is a ∼50-kb deletion (DI12) identified in ancient genomes from the first and second plague pandemics including the *mgtB* and *mgtC* loci, involved in *Y. pestis* intracellular survival when infecting macrophages^6,43^, which appears to have also independently emerged in several third pandemic 1. ORI genomes (**Fig. 5a**). We also observed uneven prevalences of the DI between lineages; while many are observed in 1.ORI, they rarely affect >1% of total isolates within this lineage (Extended Data Table 1). Conversely, deletions are highly prevalent in African 1.ANT strains, notably within the 1.ANT3 clade in which up to 3.6% of the chromosome is absent in most isolates (>80%) due to the accumulation of five DI (**Fig. 5a**, Extended Data Fig. 7, Supplementary Fig. 9).

**Figure 5.**
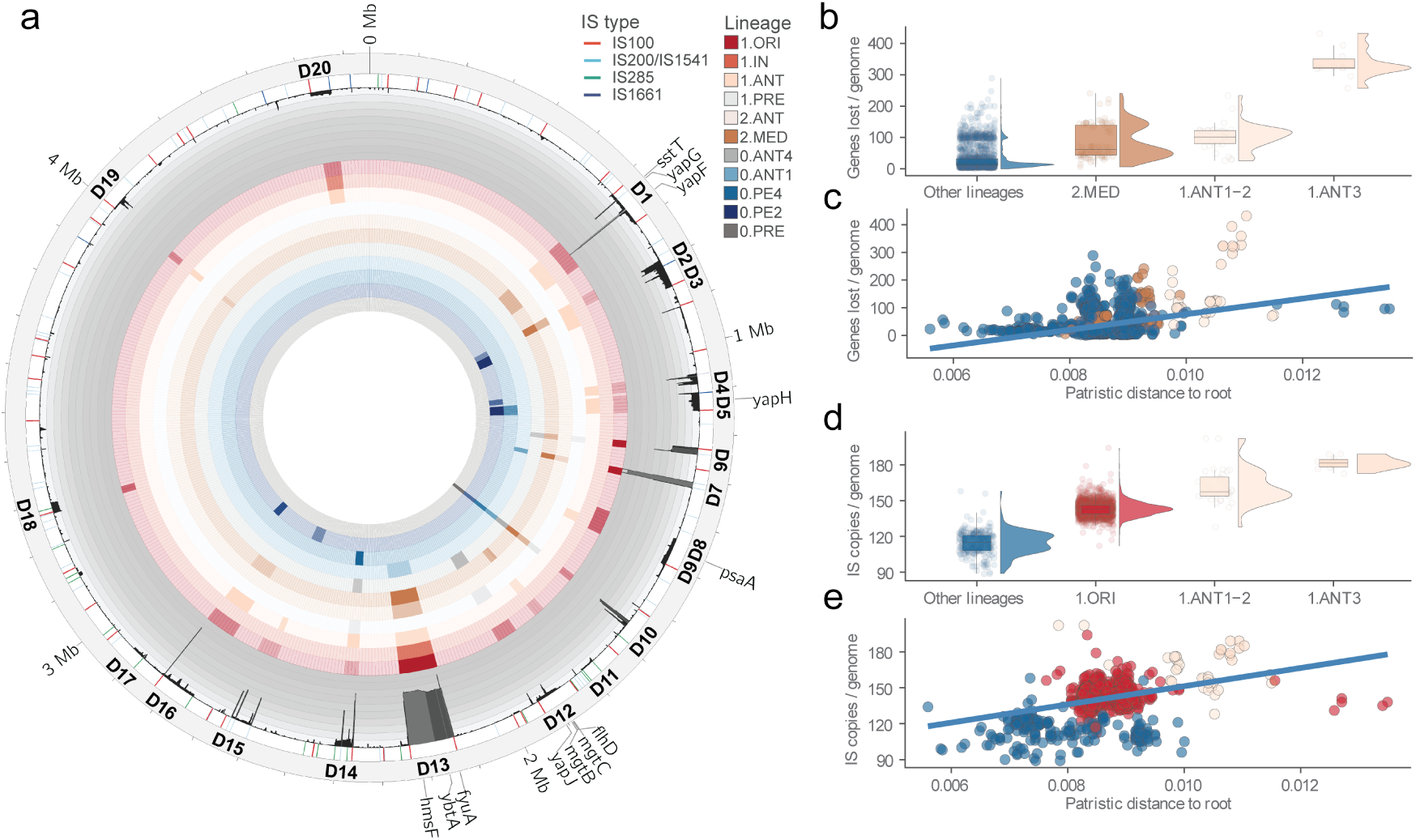
Cartography of large deletions in *Y. pestis* genomes and parallel evolutionary trajectories of gene loss and IS expansion. **a**, Distribution of the 20 deletion islands (DI) identified along *Y. pestis* chromosome (see Extended Data Table 1 and Supplementary Table 13 for the detailed coordinates and gene content of each DI). Each ring represents, from the outermost to the innermost: i) reference chromosome CO92 chromosome with the indicated DI; ii) coordinates of different IS element families present in the reference chromosome, as most identified DI appear flanked by IS; iii) histogram of the sequencing coverage estimated by chromosomal windows of 2 kb across the complete dataset of 2,806 genomes. Drops are proportional to the number of sequences without sequencing coverage, and the total height of the histogram represents 1,000 genomes; iii) Heatmaps of the prevalence of each DI across different *Y. pestis* lineages (one ring each, ordered following the legend), with darker color indicating higher prevalence within the lineage. Refer to Supplementary Fig. 9 for the complete heatmap of prevalences across different clades. **b**, Distribution of the total number of genes lost per genome across 2,128 modern genomes of different lineages. 1.ANT genomes show higher gene loss rates compared to all other lineages (Wilcoxon-Mann-Whitney test, *p*<0.001). See Supplementary Table 12 recapitulating the total gene loss in each genome. **c**, Gene loss trajectory in *Y. pestis* lineages. Scatterplot of total genes lost per genome and the genetic distance to root (patristic root-to-tip distance) estimated from the phylogenetic tree. Dots are colored by lineage as in panel b. The fitted line shows the linear regression of IS copy number against root-to-tip distance per genome, with a single slope fit to all lineages (correlation assessed using a permutation test, *p*<0.001, see Methods). **d**, Distribution of total IS copies per genome across 2,128 modern genomes. IS burden is significantly higher in 1.ANT and 1.ORI compared to all other lineages (Wilcoxon-Mann-Whitney test, *p*<0.001). For the detailed prevalences and IS families in each genome, see Supplementary Table 15. **e**, IS expansion trajectory in *Y. pestis* lineages. Scatterplot of IS copy number per genome and its corresponding patristic root-to-tip distance, with dots colored by lineage as in panel d. The fitted line corresponds to the linear regression between the IS copy number in each genome and its genetic distance to the root (correlation assessed using a permutation test, *p*<0.001).

To better characterize the extent of gene loss in *Y. pestis* lineages, we quantified the total genes lost or inactivated in modern genomes (Methods). Our results indicate that deletions are the most common mechanism driving gene loss in *Y. pestis* (Supplementary Fig. 10). The analysis confirmed that African 1.ANT genomes significantly show the highest gene loss rates (Wilcoxon-Mann-Whitney test, *p*<0.001), followed by 2.MED and 0.PE4 lineages (**Fig. 5b**, Supplementary Fig. 11, Supplementary Table 12). Moreover, we found a positive correlation between the genetic distance to the root of the phylogeny (root-to-tip distance) and the total genes lost in each genome (Permutation test, *p*<0.001, **Fig. 5c**), indicating that genome degradation follows the increased diversification of some *Y. pestis* lineages.

Our analysis also revealed that most DI appear to be flanked by insertion sequences (IS) elements (**Fig. 5a**, Supplementary Table 13), which are highly prevalent in *Y. pestis* and were previously associated with the loss of genomic regions and gene interruptions due to their recombination and transposition activity^12,13^. We thus speculated that IS activity may have contributed to the gene loss trend that we observed and the particularly higher gene loss rate in 1.ANT genomes. To explore the dynamics of IS in *Y. pestis* evolution, we analysed the frequency and distribution of six IS types commonly present in completed genomes of *Y. pestis*: *IS1541/IS200*, *IS100*, *IS285*, *IS1661*, *ISYpe1* and *ISYps7* (Supplementary Table 14). We then identified chromosomal insertion sites for these IS in short-read data of 2,128 modern *Y. pestis* genomes (see Methods). Significant differences in chromosomal IS counts were observed between lineages (89-202 copies, median 142), with African 1.ANT genomes and particularly 1.ANT3 genomes carrying the highest number of IS insertions (Wilcoxon-Mann-Whitney test, *p*<0.001, **Fig. 5d**, Supplementary Fig. 11, Supplementary Table 15), especially through the expansion of *IS100* copies within the lineage (Extended Data Fig. 8). Accordingly, the distribution of IS insertions within the *Y. pestis* chromosome clustered genomes following the population structure defined by the species phylogeny (Extended Data Fig. 9), indicating that IS insertions and their prevalence are a marker of lineage diversification. Similar to gene loss dynamics, a positive correlation was observed between the total IS copies and the root-to-tip distance for each genome (Permutation test, *p*<0.001, **Fig. 5e**, Supplementary Fig. 12), drawing parallel trajectories between IS expansion and gene loss in increasingly diverse *Y. pestis* lineages.

Such expansion of gene loss and increased IS burden has likely led to metabolic streamlining in some lineages, particularly in 1.ANT as it has lost or inactivated hundreds of genes (median 323 in 1.ANT3 genomes). From a functional standpoint, the DI identified in our work (**Fig. 5a** and Supplementary Table 13) include several bacterial surface and membrane transport factors known to contribute to *Y. pestis* virulence such as YapJ and PsaA adhesins^37,44^, as well as RbsA transporter^45^, raising questions regarding the impact of such deletions on *Y. pestis* infection biology and their evolutionary implications. We also detected 19 chromosomal loci displaying an enrichment of IS integrations within their sequence, likely causing their loss of function (Supplementary Fig. 13, Supplementary Table 16). Four of these genes (*rnd*, *cytR*, *pldA, YPO0623*) also appear within the genes with higher mutation density identified above, in which at least half of their mutations are nonsense (Supplementary Fig. 5) and have also been shown to accumulate multiple small insertions/deletions (indels)^23^. This finding indicates that IS indeed participate in the adaptive inactivation of some functions in *Y. pestis* evolution. Altogether, our analyses show that gene loss is a major mechanism fueling *Y. pestis* genome diversification, likely associated with genomic plasticity driven by expanding IS elements in the evolving populations. This trend is observed throughout the evolution of different *Y. pestis* lineages but notably expanded in 1.ANT genomes, which might have contributed to the local adaption and persistence of this lineage in Central-Eastern Africa.

## Discussion

Previous population genomics studies of *Y. pestis* have mainly characterized the diversity of lineages within vast regions across Eurasia^5,15^ and have retraced its evolution and spread in historical pandemics and outbreaks with ancient DNA^6,8,16^, while diversity in endemic regions of continental Africa has been notably missing from this global picture. Here, by analyzing a rich collection of African and worldwide isolates, we nearly doubled the amount of available genomic data of *Y. pestis*, providing a more complete outline of the species global population structure and revealing key mechanisms of genome evolution driving strain-to-strain differences. The absence of antimicrobial resistance determinants within the newly sequenced isolates suggests that the emergence of resistant *Y. pestis* still represents a rare event. The global genomic dataset generated in this study will also be a key resource for the monitoring and genomic epidemiology of plague.

Our analysis illuminates the diversity of *Y. pestis* in Africa, the only non-Eurasian continent with several co-existing lineages, where the 1.ANT lineage has undergone significant genetic diversification. We also provide evidence for the expansion of 1.ANT lineage out of Africa, as shown by a 1.ANT isolate from a Yemen outbreak, revealing a broader geographic range than previously recognized. From our molecular dating analyses, we generated in-depth estimations of 1.ANT lineage emergence and diversification, retracing the epidemic history of plague in Africa since approximately the late 17^th^ century and presenting a plausible scenario of its early introduction and dispersal patterns around the Great Lakes region and, later, the Swahili Coast. By integrating historical research and genomic information, we provide evidence for the prolonged presence of plague in Africa at least two centuries before the third plague pandemic reached the African coasts in the late 19^th^ century. The later, independent establishment of 1. MED1 and the newly described 1.ORI5 clade into continental Africa unambiguously links plague in this continent to Asian foci as their source^11,24,25^. We expect that the present historical-genomic synthesis will serve as a starting framework from which future research on the epidemiological history of plague in Africa will arise.

The global evolution of *Y. pestis* is characterized by recurrent dispersal events into new areas within and outside of Eurasia, followed by its long-term circulation in different animal reservoirs^2–4^. In this study, we also sought to characterize different genomic variation patterns emerging during the diversification of *Y. pestis* and its persistence in different endemic regions. We approached this question by exploring two levels of genomic change: 1) adaptive signatures within genes and 2) gene content variation between strains. Detecting adaptive signatures in genetically monomorphic pathogens like *Y. pestis* is challenging due to their relatively recent emergence and limited diversity^46^. For these reasons, a previous study conducted on just a few hundred genomes could not find unifying functions in the few genes showing potential adaptive signatures^5^. Conversely, we scrutinized variants from thousands of modern *Y. pestis* genomes from diverse genetic and geographic backgrounds and identified potential adaptive signals mainly within regulatory factors known to contribute to virulence and biofilm formation in pathogenic bacteria^34,36,37,47^. As biofilm production rates and virulence are two essential aspects determining *Y. pestis* ability to follow a vector-borne transmission by fleas and infect different mammal hosts^33^, it is likely that the variants in the identified genes contribute to the adaptation and persistence of *Y. pestis* in endemic foci. The placement of these mutations across the phylogeny may represent adaptations to fluctuating conditions the pathogen encounters within its ecological niche, in agreement with recent results showing the transient emergence of genetic variants modulating biofilm rates in *Y. pestis*^19^ and a higher variation in few genes related to stress response^23^.

In particular, the observed over-representation of mutations at DNA-binding residues of RovA suggests that adaptive pressures have shaped its diversification. RovA is a transcription factor which plays a pivotal role in the intricate interplay between the expression of virulence and biofilm genes in *Y. pestis* together with other factors like RovM, HmsT and PhoP/PhoQ^33,37^. From our *in-silico* analysis we can only speculate on the possible impact of the identified variants on RovA activity, although some of them have previously been shown to reduce its transcriptional activity^40^. RovA impairment would provide higher transmission rates in the flea while moderating virulence within the host^33^, a trade-off that may be advantageous in some host/vector population dynamics. Given the global regulatory roles of RovA and other identified factors with adaptive signatures, the phenotypic effect of their genetic variants on *Y. pestis* pathogenicity and transmission is worthy to be experimentally investigated.

Apart from point mutations, this study uniquely captures the dynamics of gene loss and IS expansion as *Y. pestis* lineages diversify. Our findings predict a genome degradation trajectory in the evolution of several *Y. pestis* lineages, a trend likely driven by genetic drift, weaker purifying selection^46^ and the parallel expansion of IS, which make the genome more flexible and prone to deletions through recombination between IS copies^13^. Proliferation of IS elements appears to be mainly due to the activity of *IS100*, which is present in all three plasmids commonly present in *Y. pestis*^12^ and thus their expansion may be plasmid-driven. Once large deletions occur during the emergence and evolution of lineages, they are unlikely to be regained due to the limited probability of recombination between geographically distant strains. Although ours and previous analyses demonstrate the acquisition of new plasmids in some *Y. pestis* clades^15,41,48^, these events are of low prevalence. By contrast, the extent of gene loss, which involves hundreds of genes across different clades, suggests that such events may have a greater impact on *Y. pestis* genome evolution compared to gene gain. Large deletions and the specific IS burden thus serve as genetic markers, distinguishing different lineages and clades in terms of gene content. This finding complements recent observations that small indels mirror the phylogenetic relationships of *Y. pestis* genomes^23^.

Genome degradation is a hallmark of pathogens emerging from evolutionary bottlenecks and adapting to a narrow ecological niche^49^. In the case of *Y. pestis*, gene loss is one of the major processes that guided its adaptative emergence as a vector-borne pathogen when it evolved from its enteropathogenic ancestor, *Y. pseudotuberculosis*^1,12^. Our analysis identifies extensive gene loss and IS burden in African 1.ANT genomes and the convergent deletion of the several genomic islands between lineages, suggesting that loss of certain metabolic functions may be advantageous under certain conditions and ecological niches that remain to be characterized^23^. Gene loss events are known to contribute to niche restriction by reducing metabolic flexibility^13,49^. Therefore, one possibility may be that genome degradation is mediating *Y. pestis* specialization to particular animal host species in endemic foci. Host specificities have indeed been observed in some *Y. pestis* populations, including Asian 0.PE4^5,41,50^, which also shows high gene loss rates. By losing certain metabolic capabilities, metabolic specialization would contribute to the maintenance of host-adapted lineages in endemic foci, a process that may have facilitated the long-term persistence of the 1.ANT lineage in Africa given its extended genome degradation. The comparative analysis of metabolic pathways lost among lineages, backed by modeling and phenotypic testing, will help to elucidate the impact of gene loss on the metabolic capabilities of different *Y. pestis* lineages.

In conclusion, in this study we integrate phylogenomic and historical approaches to retrace the evolutionary history of *Y. pestis* in continental Africa and explore species-wide genome variation mechanisms, revealing two recurrent patterns in *Y. pestis* evolution: a signature of increased diversification of virulence and transmission determinants; and a genome degradation trajectory in divergent lineages, which is particularly sharp in African 1.ANT genomes. Our ability to detect these patterns was made possible only by leveraging the significant increase in the number and worldwide diversity of the *Y. pestis* genomes sequenced here, which highlight the significance and reach of our study. We anticipate that further investigations of these aspects will uncover factors that have enabled one of the most virulent pandemic pathogens to persist and reemerge multiple times over the past millennia.

## Data availability

All raw sequencing reads generated in this study have been submitted to NCBI under accession number PRJNA1137242. Additional data is available within the Supplementary Information section and Supplementary Tables.

## Code availability

All code used to generate the figures and statistical analysis of this study and the XML file used for the time-scaled phylogenetic inference in BEAST2 are available at https://github.com/guillemmasfiol/GlobalPlague. The global phylogeny with available metadata have been imported into a project in MicroReact accessible at https://microreact.org/project/qDejD9oBnnfbhDvE2PgHfV-global-phylogeny-of-yersinia-pestis-species.

### Acknowledgments

We thank François-Xavier Weill, Eduardo Rocha, Julian Parkhill, Sandra Reuter, Kate Baker, and Hendrik Poinar for critical comments during the development of this project; all other members of the Yersinia Unit the Institut Pasteur for helpful discussions throughout the course of this study; Sylvie Brémont for the assistance in the recovery of isolates of old *Y. pestis* collections. This work stems from the collaborative heritage of the historical collection of isolates that have been recovered and sent from diverse sources over the past century by many different actors, to which we are grateful. We would like to acknowledge the GPU lab of the INCEPTION program and the HPC Core Facility of Institut Pasteur for providing access and support to use computing resources.

## Author contributions

G.M.F., P.S., N.R. and J.P.C conceived and led the investigation. G.M.F., G.B., D.M., F.L., A.A.V., P.S., N.R. and J.P.C designed the study. G.M.F., A.K., C.B. and R.B. performed the laboratory work. E.C., M.C., M.R. and J.C.S. provided bacterial isolates. G.M.F., D.M., F.L., A.A.V. and C.S. performed the genomic data analysis. G.M.F. and S.D. performed the molecular dating analysis. G.B. developed the structural analysis on RovA variants. P.S. assembled, analysed and translated the historical and context information. G.M.F., G.B., D.M., F.L., A.A.V., P.S., N.R. and J.P.C aided in interpreting the results. G.M.F., P.S., N.R. and J.P.C. wrote the manuscript with contributions from all co-authors.

## Methods

### Bacterial isolates and genome sequencing

A global collection of 1,124 *Y. pestis* isolates was collected by different teams and Pasteur Institutes around the world between 1899 and 2013, in the context of human outbreaks and of plague surveillance tasks (Supplementary Table 1). All manipulations of live *Y. pestis* isolates were performed in biosafety level 3 laboratory conditions at the Institut Pasteur in Paris. Sequencing libraries were prepared following the protocol of the Nextera DNA Flex library preparation kit (Illumina). Whole-genome sequencing reads of the isolates were obtained using an Illumina MiSeq platform, generating pair-end sets of 2×300 bp lengths and a median of more than 2M reads per library.

### Sequence analysis and *de novo* assembly

The sequencing reads of the 1,124 isolates from the Pasteur collection were added to a larger genomic dataset that included most of the available *Y. pestis* samples available in public repositories (n=1,682, Supplementary Table 2). This dataset included ancient DNA metagenomes from different centuries that have been analyzed in recent studies (n=136), as well as genomic assemblies (n=401) from modern isolates downloaded public repositories (NCBI, EnteroBase^1^). For the 1,124 newly sequenced *Y. pestis* isolates, *de novo* assemblies were generated using the fd2dna 21.06 workflow (https://gitlab.pasteur.fr/ GIPhy/fq2dna), which was run using default parameters and includes the steps described below. Correction of reads, clipping and trimming of adapter sequences and non-confident calls was performed using AlienTrimmer 0.4.0^2^, and redundant or over-represented reads were identified and reduced using the khmer tool package 1.3^3^. A minimum threshold of 12x was used in the pre-processing steps, and sequencing errors were identified and corrected by Musket 1.1^4^. Preprocessed reads were finally *de novo* assembled into contigs or scaffold sequences using SPAdes 3.1.2.0^5^. Prediction of antimicrobial resistance determinants sequences was performed by querying all the assembled genomes against ResFinder 4.0 server database^6^.

### Phylogenetic analyses

To process the large dataset of samples in this study, we implemented a workflow available at a Git-Lab repository: https://gitlab.pasteur.fr/metapaleo/amphy. This workflow allowed us to integrate raw sequencing data from modern *Y. pestis* genomes and ancient DNA metagenomes, together with samples for which only contigs were available. For contigs, reads were simulated using wgsim function in SAMtools 1.13^7^. Reads for all samples were trimmed for adapter sequences using AdapterRemoval 2.3.1^8^; only bases with a quality of 2 or less were removed and reads larger than 30 bp were retained. Trimmed reads for paired-end data were collapsed and merged and mapped against *Y. pestis* CO92 reference genome containing the chromosome (NC_003143.1) and plasmids (NC_003131.1, NC_003134.1, NC_003132.1) using BWA mem 0.7.17^9^. Aligned reads were processed using SAMtools 1.13, discarding reads with an overall mapping quality lower than 30. Duplicated reads were detected and removed using MarkDuplicates from Picard-Tools 2.23.8^10^. Bam files corresponding to libraries from the same sample were merged with SAMtools 1.13 before realignment using GATK 4.1.9^11^. For ancient DNA samples, further steps were necessary to account for DNA damage. Prior to genotyping with GATK, ancient DNA bam files were rescaled using mapDamage 2.2.1^12^ to account for misincorporations of deaminated cytosines (i.e.: masking the C>T and G>A mutations in agreement with cytosine deamination at the end of the fragments present in ancient DNA^13^). Genotypes were called from the processed reads aligned on *Y. pestis* CO92 genome using GATK HaplotypeCaller. Variant sites were filtered for a minimum 4x depth for ancient DNA samples or 10x for modern ones, and maximum 1000x depth. A minimum base quality score of 20 was also used, and calls were filtered for allelic balance of 0.1. Indels were removed, while deletions were replaced by “N” and resulting VCF files were transformed into consensus fasta sequences. Coding sequences were extracted from the consensus using gene annotations from the reference genome and only genes with less than 10% of missing calls (“N”) for each sample were kept in a first step, while only genes kept in at least 50% of samples were retained for further steps. Gene consensus sequences passing the filters mentioned above for each file were then aligned and used for downstream phylogenetic analyses. A total of 2,806 *Y. pestis* genomes passing the previously mentioned filters were retained (Supplementary Table 2). To analyze the population structure of *Y. pestis* species including all the retained samples, a phylogeny was reconstructed. For this phylogenetic inference, a distance matrix was first computed from consensus sequences using Goalign 0.3.6^14^ with the F81 model. A distance tree was reconstructed from the matrix of distances using fastME 2.1.6^15^, which was then used as starting tree for a maximum-likelihood phylogenetic inference with RAxML-NG 1.0.2^16^ using a GTR+Gamma substitution model. Finally, 100 Bootstrap distance trees were computed using fastME 2.1.6^15^ to estimate branch supports. The phylogenetic tree visualization displayed in **Fig. 1** was generated using GrapeTree 1.5.0^17^. The inferred phylogeny and available metadata for the 2,806 *Y. pestis* genomes have been integrated into a project in MicroReact, which is accessible via the following link: https://microreact.org/project/qDejD9oBnnfbhDvE2PgHfV-global-phylogeny-of-yersinia-pestis-species.

### Estimation of genetic diversity statistics

Average nucleotide diversity (*π*) within modern lineages was computed by 10-kb windows from variant tables (VCF) of 2,272 modern genomes using Pixy 1.2.6, which considers missing genotypes to produce unbiased estimates^18^. Pairwise SNV distances between genomes were computed from the filtered variants identified in the phylogenetic analysis step using harrietr R-package^19^ and counts were grouped for each lineage. Only variable positions without partial deletion were used in SNV distance calculations.

### Time-calibrated phylogenetic analysis

The time-scaled phylogenetic tree (Supplementary Fig. 4) was reconstructed in BEAST2 2.6.0^20^. We used a subset of 200 *Y. pestis* genomes representative of all known major lineages and clades (Supplementary Table 4) including new genomes obtained in this study to date their divergence, together with the sequence of *Y. pseudotuberculosis* IP32953 used as outgroup to root the tree. The aligned data used to build the maximum-likelihood tree mentioned above was partitioned following codon positions and based on previous experiments^21,22^, a constant population size coalescent and an unlinked GTR substitution model with four gamma rate categories were used. A relaxed log normal clock was assumed for modelling the mutation rates within branches, as previous analyses have found strong evidence for the adequacy of loose clocks for modelling *Y. pestis* evolution^23,24^, and a uniform prior distribution ranging between 2 × 10^−6^ and 2 × 10^−8^ substitutions per site per year was used for the molecular clock model. For the BEAST2 analysis, 7 independent Markov Chain Monte Carlo (MCMC) were run with 200 million states each, sampling every 1000 states and first 10% states of each run were removed as burn-in. The effective sample sizes (ESS) of all estimated parameters were all >200. Trees of the combined runs were summarized using TreeAnnotator^25^ to generate a maximum clade credibility (MCC) tree annotated based on median heights.

### Identification of adaptive signatures in *Y. pestis* genomes

To identify high-quality variants within genes, sequences from modern genomes (n=2,700 genomes, and *Y. pseudotuberculosis* IP32953 genome as outgroup) were mapped against the CO92 reference chromosome using RedDog pipeline 1beta.11^26^, keeping only positions which are covered in at least 90% of samples (-c flag). Variants within repetitive regions (IS, heteropolymeric tracts…) were filtered out (hardfilter flag) using previous annotations on the reference chromosome NC_003143.1^27^. A maximum-likelihood phylogenetic tree was reconstructed from the variant alignment using IQ-TREE 2.2.2.2^28^ with best-fitted substitution model identified using ModelFinder^29^. The statistical approach to identifying genes with higher mutation density in *Y. pestis* genomes is described in Supplementary Information 3. Pairwise Pearson correlation coefficients between pairs of the 60 identified genes displaying higher mutation densities were calculated based on mutation occurrences across genomes and plotted using corrplot R-package.

To investigate the extension of the observed diversity in the identified genes with higher diversification that is due to mutations in African genomes, mutations in these 60 genes were grouped by gene, and for each we identified the samples that harbored at least one mutation. Using the available metadata, samples were classified into two categories based on geographic origin (African and all other geographic regions). For each gene, we calculated the total number of samples presenting mutations, as well as the number of samples within the African and non-African groups. Mutation counts were subsequently normalized to obtain proportions, reflecting the relative frequency of mutated alleles in African versus non-African strains.

As signatures of parallel evolution, homoplastic mutation events were inferred by mapping the variants back to the rooted phylogenetic tree using SNPPar 1.2^30^. For this analysis, a strategy developed by Edwards et al.^30^ was followed, in which mutation events affecting the root node and the outgroup (*Y. pseudotuberculosis*) were removed, and only unambiguous mutation events were kept (n=6,416). Genes overlapping with repetitive regions were excluded, and the total homoplastic mutation events were calculated for each gene (n=3,401 genes) using the script countMEbyGene.py. The total, nonsynonymous and synonymous mutation events per gene are described in Supplementary Table 7. Protein-coding genes were finally classified into different COG categories using eggNOG-mapper 2.1.2^31^. All code used to perform the analyses mentioned above are available at https://github.com/guillemmasfiol/GlobalPlague.

### Mapping of variants on RovA structure

Crystal structure of RovA from *Y. pseudotuberculosis* in complex with a RovA promoter fragment^32^ (PDB id: 4AIJ) was used to compute the structural features related to the identified mutations. Distances between all residues (including residues presenting mutations) and DNA were measured using python PyMOL scripts^33^. For each amino acid residue, all inter-atomic distances were computed and aggregated to obtain one distance per residue using the minimum distance. Amino acid residues depicted as interacting with the DNA fragment were designated using an upper distance threshold of 4Å. Molecular structures were rendered using PyMOL 3.0.3.

### Detection of acquired genetic mobile elements

MOB-suite 3.0.1^34^ was used with default parameters to query all *Y. pestis* genome assemblies against its database in order to identify and reconstruct putative plasmid sequences present within the genomes. Plasmid sequences identified were named based on the previous description of the replicons, while new plasmids without homology to previously described plasmids are named according to the name of the isolate in which they have been identified (for pIP2235H, pIP1930H, pIP1535H, pIP1168H). ICEfinder^35^ tool was used to identify the putative integrative and conjugative element (ICE) present within multiple assemblies. The detected sequences were extracted from the assemblies and annotated using Prokka 1.14.5^36^ and Bakta 1.5.1^37^. All identified genetic mobile elements within the assemblies and their features are listed in Supplementary Tables 10 and 11.

### Gene loss analyses

A scan of clusters of uncovered regions in the chromosome was carried out to generate a cartography of large deletions in *Y. pestis* populations. Clean reads mapped onto CO92 reference chromosome obtained from the phylogenetic inference were kept for the complete dataset of 2,806 ancient and modern genomes. Bam files with the aligned reads were used to calculate the sequencing coverages in all genomes using Bedtools 2.29.2 genomecov^38^ and then their distribution was analysed in R 4.4.0. Chromosomal regions longer than 5kb which lacked sequencing coverage in between 50 and 2,000 samples were identified and marked as deletion islands (DI). The region including a filamentous prophage YpfΦ identified in CO92 (genome coordinates 2,554,372 – 2,562,880) and other *Y. pestis* isolates was excluded from the analysis as its presence is considered an acquisition by the genomes showing coverage on its sequence^39^. The distribution of the number of genomes without sequencing coverage was computed by 2-kb chromosomal windows using bioinfo-scripts^40^ tools and the prevalence of each DI across *Y. pestis* clades were visualized using Circos 0.69-9^41^. To help the visualization of the identified DI, non-contiguous chromosomal windows uncovered in more than 400 genomes were filtered out, as they globally corresponded to insertion sequences. The presence of each DI for each genome was plotted next to the complete phylogeny using ggtree R-package^42^.

To investigate the extent and dynamics of gene loss in *Y. pestis* genomes, average sequencing depths for the reference chromosome (NC_003143.1) were calculated using RedDog pipeline and samples with average sequencing depths of at least 50x were retained (2,128 genomes) to ensure that uncovered regions represented true deletions. Chromosomal genes without sequencing coverage were considered as deleted and intragenic nonsense mutations were identified for each genome from the RedDog output. IS insertions falling within genic coordinates were identified from the output table from ISMapper (see the following section on IS analyses). The number of genes affected in each of the mechanisms of gene loss was computed for each genome and their sum was calculated to obtain the burden of gene loss per genome (Supplementary Table 12). n=70 genes (from YPO_RS10485 to YPO_RS10830 in CO92 reference), part of the *pgm* locus, were excluded from the total counts to avoid sampling bias, as this unstable genomic island is known to be frequently lost when growing the isolates *in vitro*. For each genome, the patristic distance to the root of the previously generated phylogenetic tree of modern genomes were calculated using ape R-package^43^ and samples with terminal branch length higher than two standard deviations (27 genomes) were marked as outliers and excluded. The correlation between the total genes lost per genome and the root-to-tip distance of each genome was assessed using a permutation test to account for the statistical non-independence of related genomes (Supplementary Fig. 12).

### Insertion sequences analyses

IS present in *Y. pestis* CO92 reference genome were detected using ISSaga^44^ for IS sequences that are present in the curated database ISFinder^45^. IS types that were present in at least one complete copy with 80% nucleotide identity to an IS in the database were kept (*IS100*, *IS200*/*IS1541*, *IS285*, *IS1661*, *ISYpe1* and *ISYps7*). To obtain a preliminary estimation of the IS burden in the general *Y. pestis* and *Y. pseudo-tuberculosis* populations and compare both species in terms of IS, all genomes of both species marked as ‘completed’ in PATRIC database (n=62 *Y. pestis*, n=26 *Y. pseudotuberculosis*) were downloaded. BLAST+ 2.2.3 was used to search for the six previously identified IS types within the closed genomes. Nucleotide BLAST+ hits with >95% identity and >90% coverage were kept and counts per genome are reported in Supplementary Table 14. To identify IS insertions within paired-end data, we employed the strategy developed by Hawkey et al.^46^. First, to allow more precise detection of IS insertion sites on *Y. pestis* chromosome, an IS-free version of *Y. pestis* CO92 reference chromosome was generated as follows. BLAST+ hits positions on the reference chromosome for copies of the six previously identified IS types were annotated and inspected. The regions where the identified IS types copies fell were removed from the reference chromosome to generate the IS-free reference. Annotations of the CDS and gene features in the original, complete, chromosome were then transferred to the IS-free reference using RATT^47^. The IS-free reference chromosome has been deposited in GitHub: https://github.com/guillemmasfiol/GlobalPlague. ISMapper 2.0.2^48^ was then used with default parameters to detect IS insertions for the six IS types onto the IS-free CO92 reference. Raw reads of a dataset of 2,128 modern *Y. pestis* genomes with an average sequencing depth of at least 50x onto the reference chromosome were used as required by ISMapper to obtain confident calls that are strongly supported by evidence from the reads^48^. Partial hits (i.e.: IS found spanning from the end of the replicon to the start) were identified and removed.

To place detected IS insertions together with the phylogenetic context of each genome analyzed in the dataset, IS counts were plotted next to the maximum-likelihood phylogeny of the samples using ggtree^42^ R-package. The patristic genetic distance for each taxon to the root of the tree was calculated using ape R-package^43^ and samples with terminal branch length higher than two standard deviations (27 genomes) were marked as outliers and excluded from the regression. Correlation between the total IS counts per genome and the root-to-tip distance of each genome was assessed using a permutation test (Supplementary Fig. 12). To cluster the genomes following their IS insertion patterns, a Principal Component Analysis (PCA) was run on 2,566 chromosomal IS insertions coordinates detected in the genomes using FactoMineR 2.9^49^ and visualized using factoextra 1.0.7 R-packages^50^. Four outlier samples were excluded from the PCA to improve the visualization of clusters: IP619H (this study), CMCC84046^23^, El Dorado^51^ and Java 9^52^. To identify genes presenting increased rates of IS insertions in *Y. pestis* genomes, the observed number of IS insertion sites within each locus was compared to the expected value given their sequence length, assuming IS insertion events across the chromosome follow a Poisson process. Using a Bonferroni-adjusted threshold accounting for multiple testing, n=19 genes significantly deviating from the theoretical expectations were identified and the distribution of observed and expected IS insertions per gene were drawn using ggplot2 R-package^53^.

---

**Extended Data Fig. 1.**
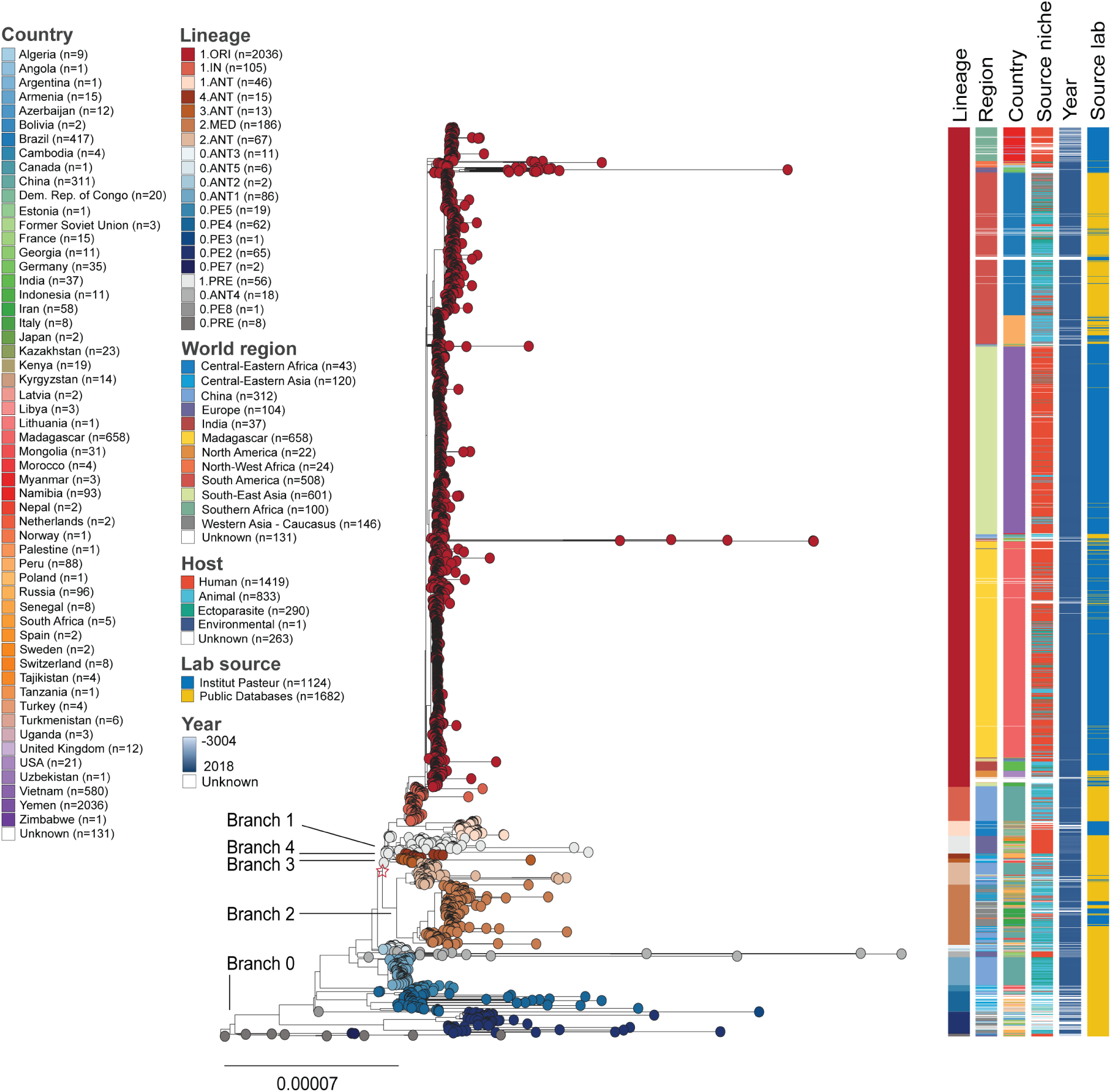
Maximum-likelihood phylogeny of the 2,806 *Y. pestis* genomes summarizing the genetic diversity of the species. The tree is reconstructed with 2,723 modern and 83 ancient *Y. pestis* genomes, based on 155,201 whole-genome polymorphic positions. Nodes defining the start of major branches (0-4) as defined in previous studies are indicated. The star highlights the node of emergence of branches 1-4, also known as “Big Bang”. Colour strips include annotations for the correspondence of genomes to 20 major lineages, world region and country of origin of the sample, isolation niche, year, and source laboratory, respectively. Colour legend of the lineages corresponds to the tree shown in Fig. 1a. Scale represents the number of substitutions per genomic position. The phylogeny and sample metadata were visualized using Microreact (https://microreact.org/project/qDejD9oBnnfbhDvE2PgHfV-global-phylogeny-of-yersinia-pestis-species).

**Extended Data Fig. 2.**
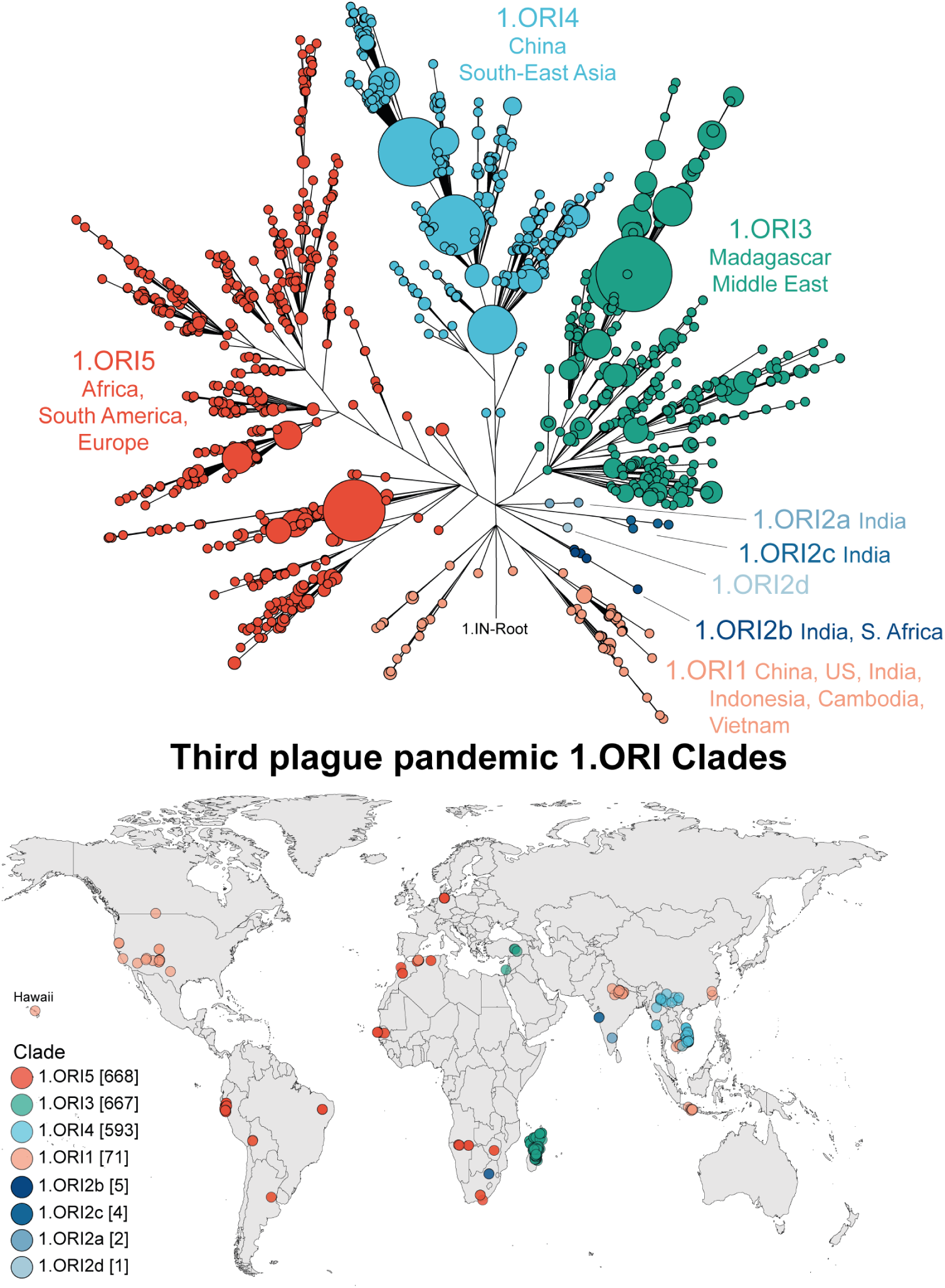
Phylogenetic relationships and geographic distribution of the 2,036 1.ORI lineage genomes associated with the third plague pandemic. Upper plot: Focus on the 1.ORI branch extracted from the global phylogenetic analysis of the species shown in Fig. 1a. The identified monophyletic clades within 1.ORI are indicated. Lower plot: Geographic distribution of 1.ORI isolates with available associated locations or country of origin.

**Extended Data Fig. 3.**
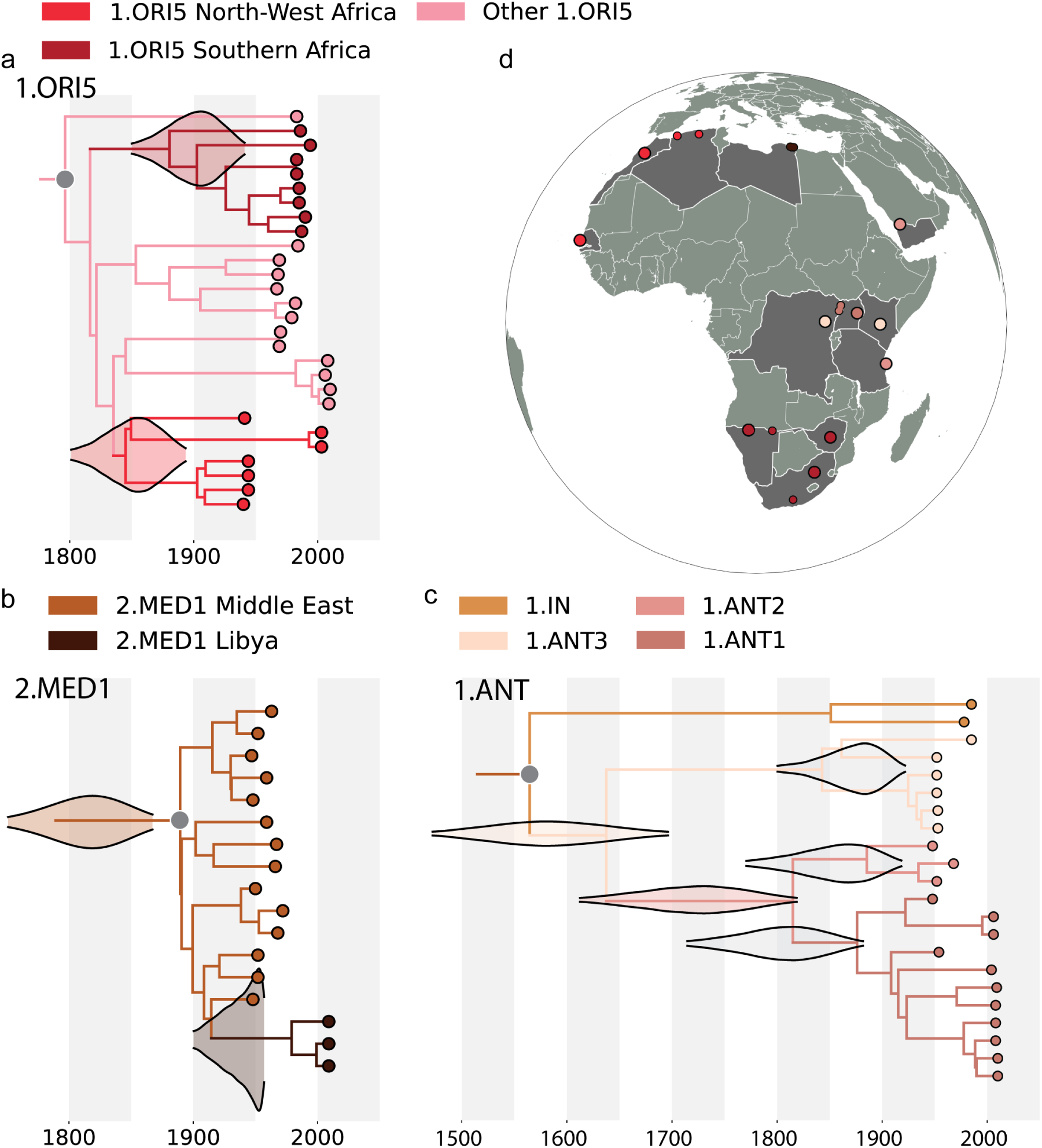
Temporal-phylogenetic relationships and geographic distribution of three different *Y. pestis* clades present in continental Africa. Panels for 1.ORI5, 2.MED1 and 1.ANT clades represent the branches extracted from the complete time-calibrated maximum clade credibility tree inferred using BEAST v2.6.6, shown in Supplementary Fig. 4. Density distributions represent the estimated posterior distributions for the divergence date of the last common ancestor of different monophyletic branches, including: **a**, the two 1.ORI5 subclades that are associated with Southern and North-West Africa; **b**, the Middle East-associated 2.MED1 subclade, which includes a branch with three genomes from Libya; and **c**, the inner clades within Central-Eastern Africa 1.ANT (bottom-right). The estimated posterior distributions for the diversification nodes within each 1.ANT clade (1.ANT3, 1.ANT2 and 1.ANT1) are displayed in grey. **d**, geographic locations for the complete set of continental African genomes of each lineage included in Fig. 1, with tips colored according to the same colors in the tree panels.

**Extended Data Fig. 4.**
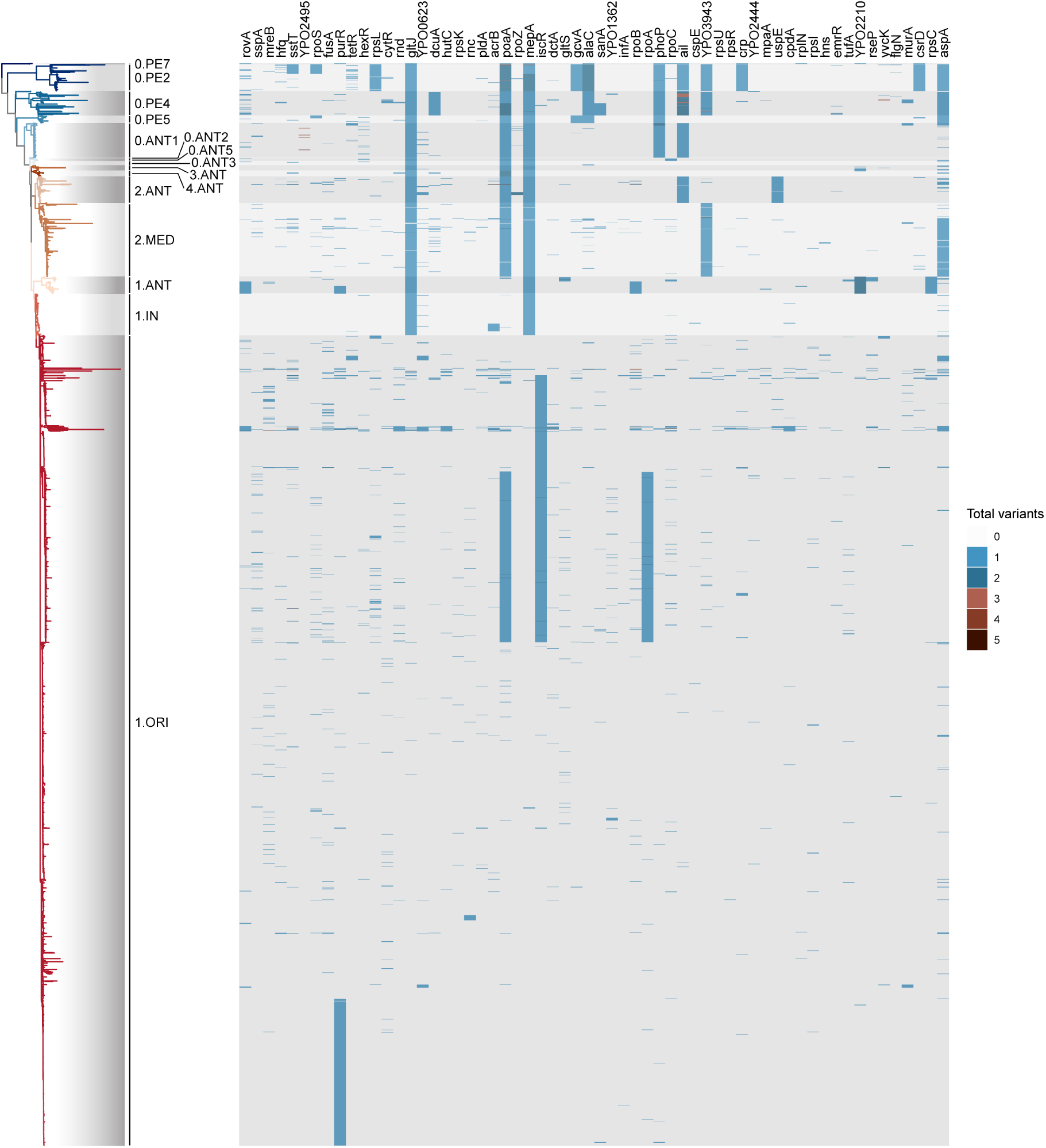
Phylogenetic distribution of total substitutions in 60 identified genes with significantly higher mutation density across 2,700 modern *Y. pestis* genomes. The tree displays the maximum-likelihood phylogeny of 2,700 *Y. pestis* isolates inferred using IQ-TREE 2.0.6. The heatmap shows the total number of substitutions in each genome for each of the 60 identified chromosomal genes showing substitution counts significantly deviating from theoretical expectations under neutral evolution.

**Extended Data Fig. 5.**
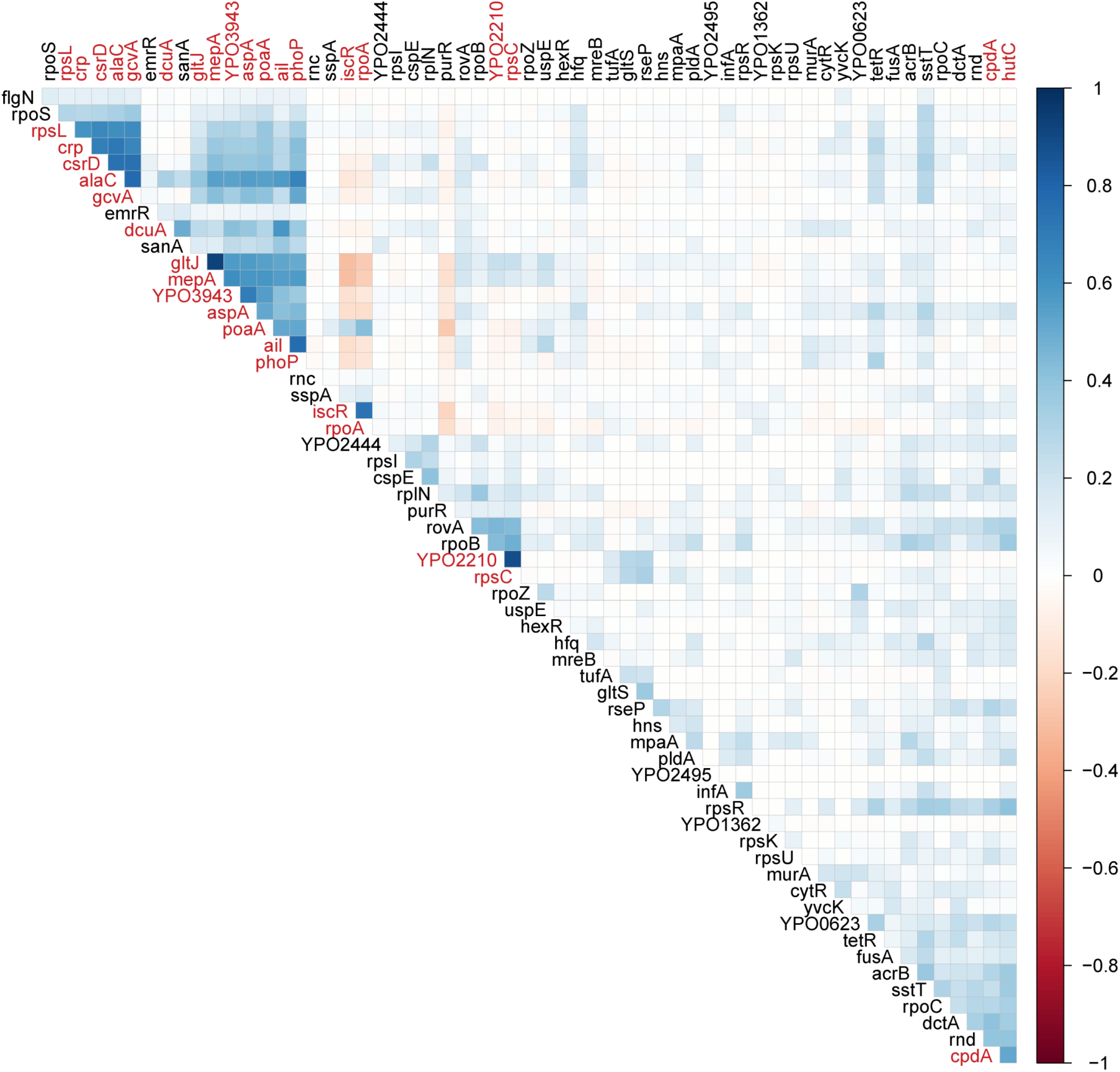
Pairwise correlations between 60 identified highly variable genes in *Y. pestis* evolution. The heatmap represents the matrix of Pearson correlation coefficients between pairs of genes, based on the total nonsynonymous mutation counts in each of the 2,700 modern genomes analyzed. A high positive correlation coefficient between a pair of genes indicates that mutations in both genes tend to occur together in the same genomes. Labels in red indicate genes with positive correlation coefficients higher than 0.5 with at least another of the identified genes.

**Extended Data Fig. 6.**
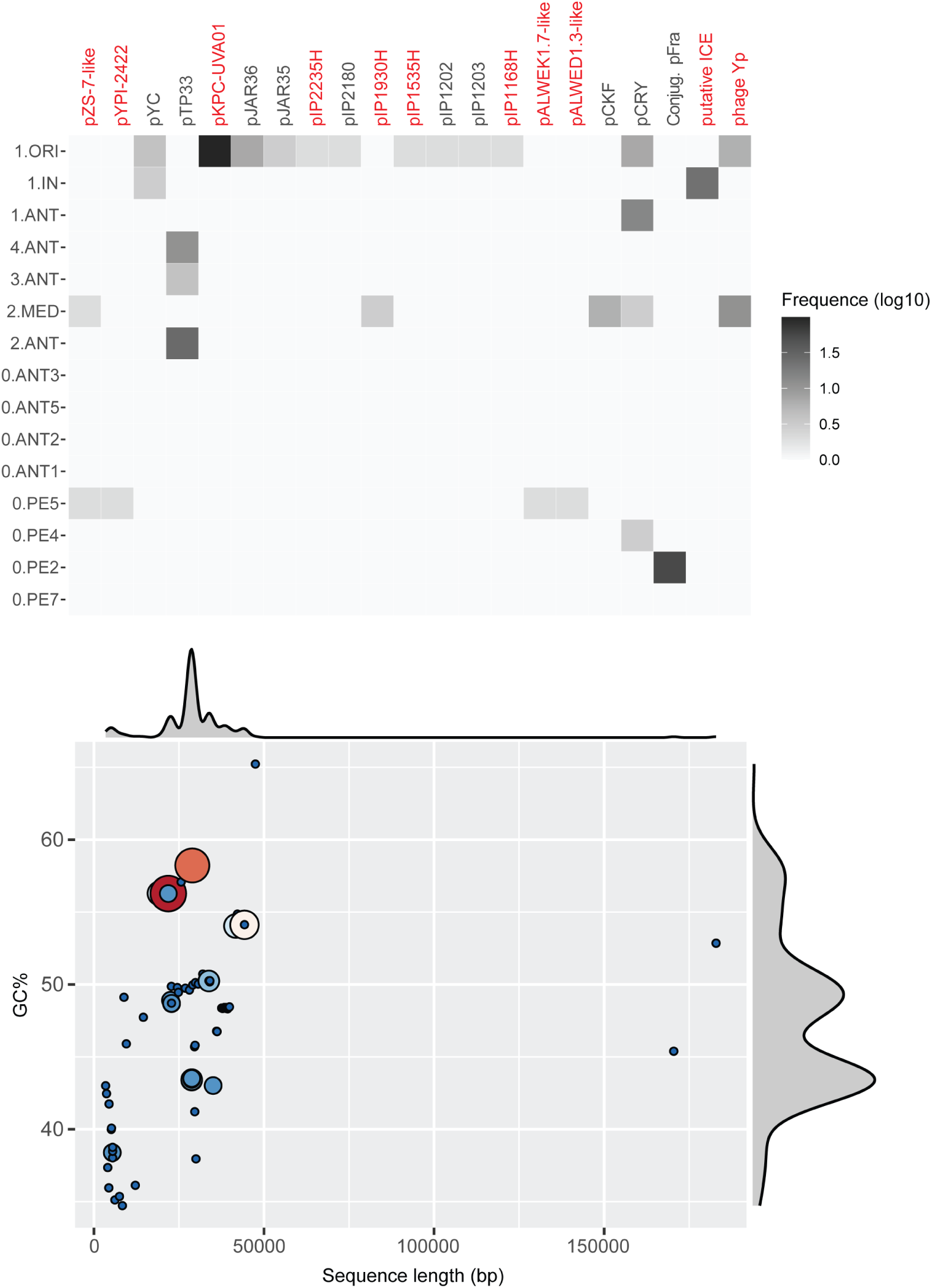
Identification of acquired genetic mobile elements in *Y. pestis* genomes. Upper panel: Heatmap of prevalence of 21 sequences variably present across different lineages, detected from the analysis of 2,692 genome assemblies. Labels highlighted in red represent sequences not previously identified in *Y. pestis* genomes. Plasmid sequences identified using MOB suite are named based on the previous description of the replicon, while new plasmids without homology to previously described plasmids are named following the name of the isolate in which they have been identified (pIP2235H, pIP1930H, pIP1535H, pIP1168H). Conjug. pFra indicates conjugative transfer-associated genes in pFra of 0.PE2 isolates; ICE indicates a putative integrative and conjugative element identified in 1.IN genomes; and phage Yp indicates a putative phage sequence identified in multiple 1.ORI and 2.MED isolates. Lower panel: Values and density distributions of average GC content and length in pairs of bases for the sequences of each of the 21 identified elements extracted from the genome assemblies.

**Extended Data Fig. 7.**
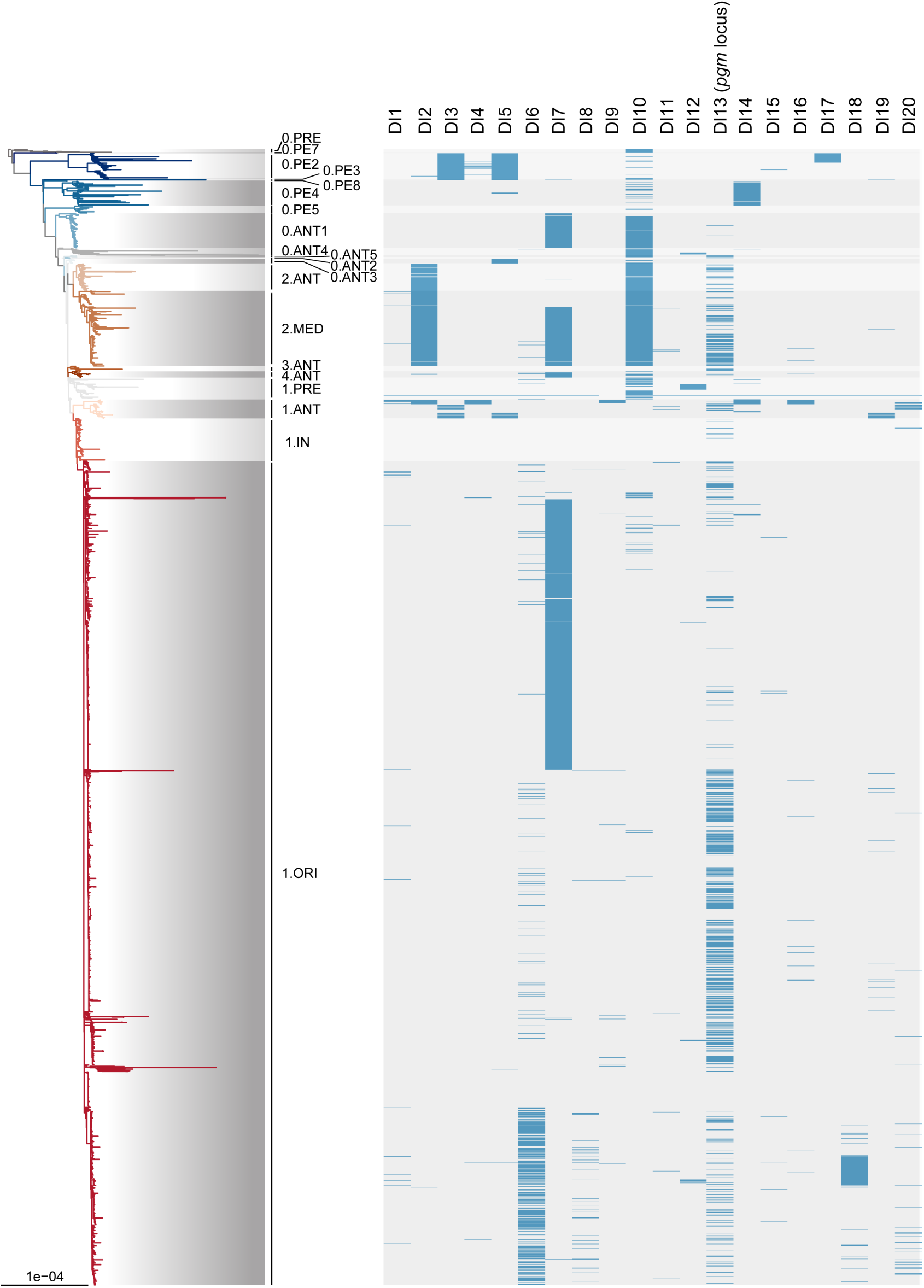
Distribution of 20 deletion islands across *Y. pestis* phylogeny. The tree represents the maximum-likelihood phylogeny of 2,806 modern and ancient genomes displayed in Fig. 1. Scale represents the number of substitutions per genomic site used to reconstruct the phylogeny. The heatmap represents the presence (blue) or absence (grey) of each of the 20 identified DI in each genome.

**Extended Data Fig. 8.**
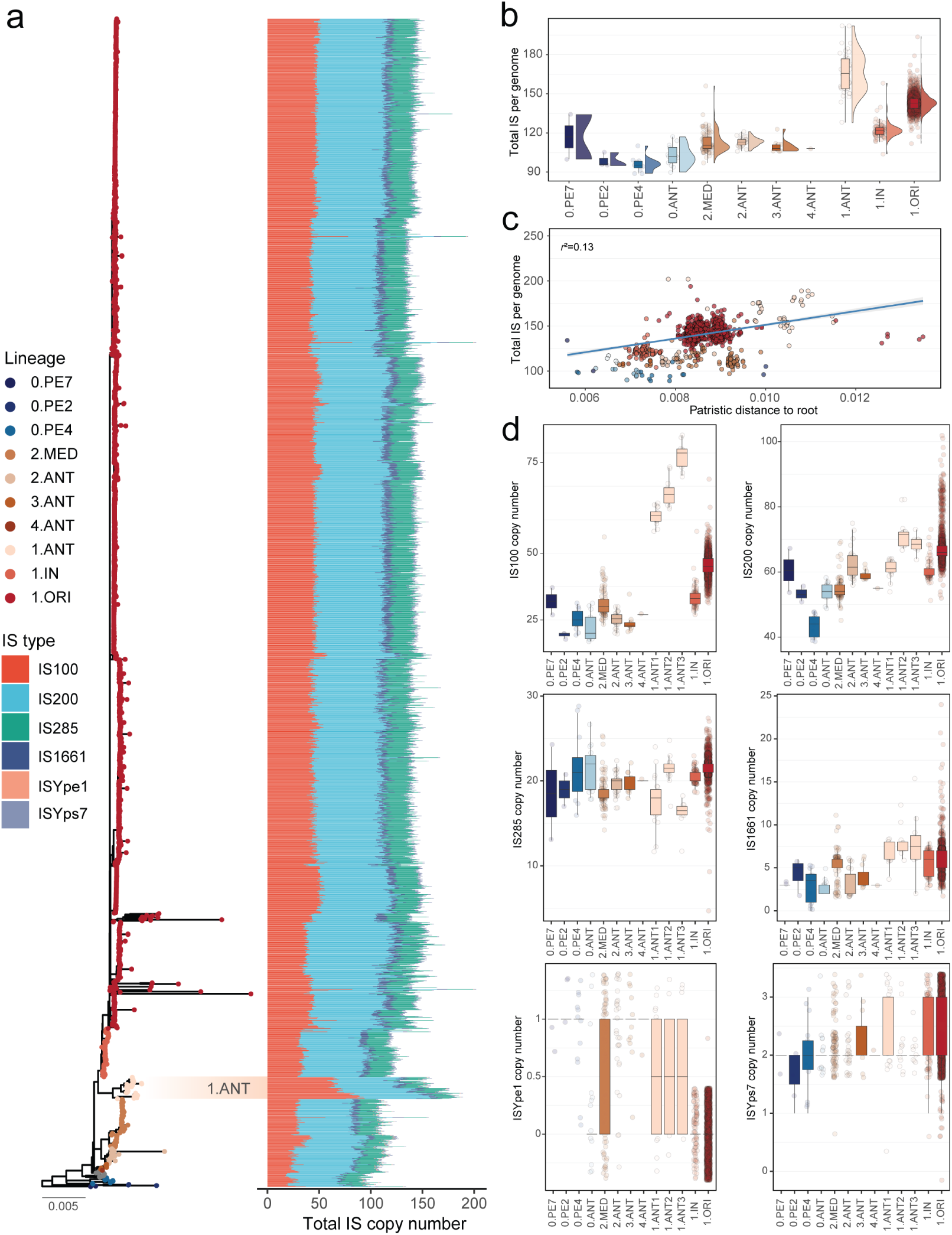
IS dynamics in *Y. pestis* evolution. **a,** IS dynamics according to *Y. pestis* population structure. Total counts of the 6 identified IS types per chromosome are represented next to the maximum-likelihood phylogeny of 2,128 genomes, highlighting the counts of 1.ANT genomes. The tree was rooted on *Y. pseudotuberculosis* IP32953 (not shown) and is ordered from most ancestral (down) to more recently evolved lineages (up). **b**, Distribution of total chromosomal IS copy number per genome across different lineages. **c**, Linear regression analysis between the total IS counts and the patristic genetic distance to the root of the phylogeny for each genome shown in panel a. Pearson correlation coefficient is shown, and significance of the regression was assessed using a permutation test (*p*<0.001). Dots are colored according to lineages correspondences of the tree. **d**, Boxplots showing the distribution of the copy number for the 6 IS types across modern lineages included in the analysis in order to identify the IS types that have expanded the most through *Y. pestis* evolution.

**Extended Data Fig. 9.**
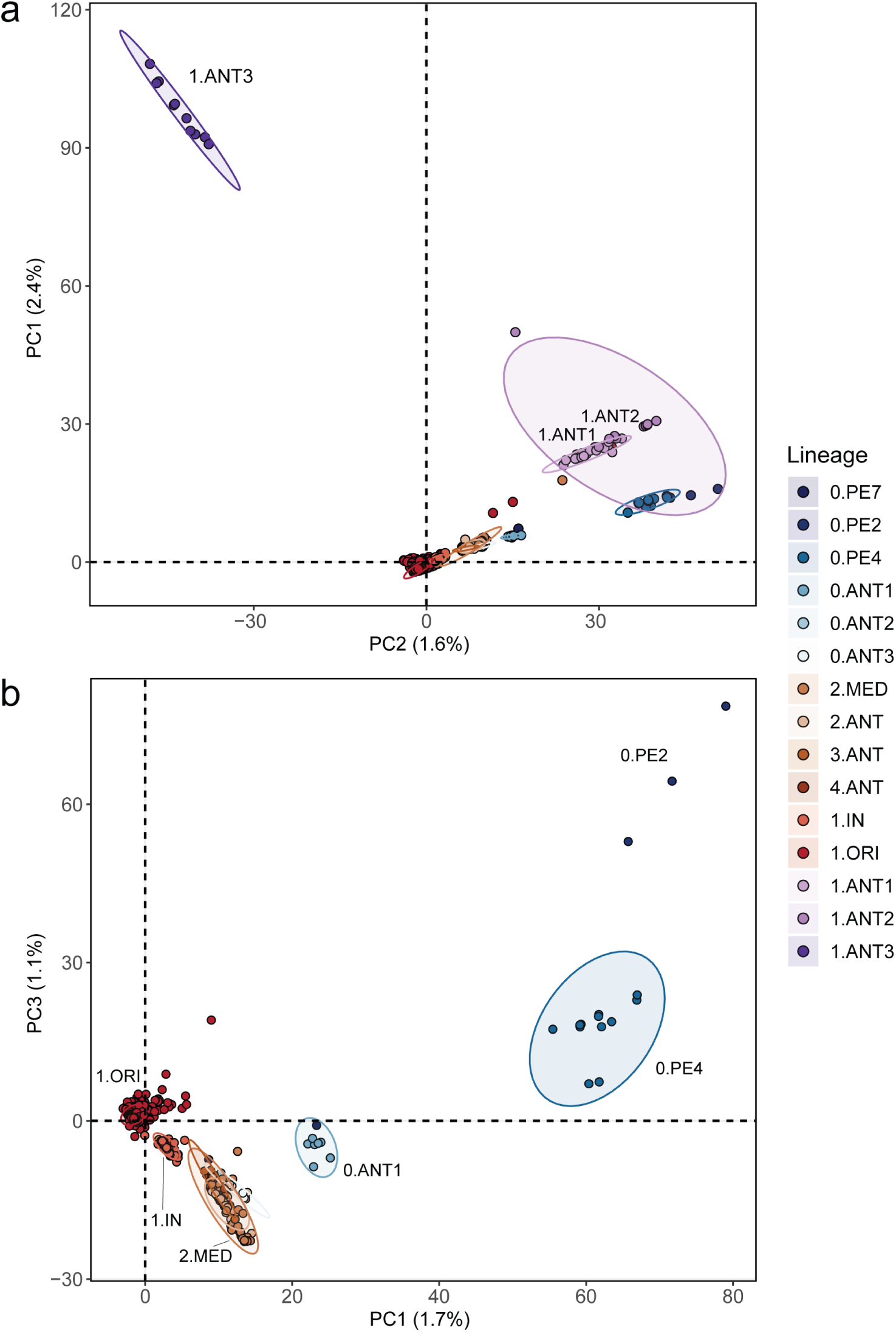
Principal component analysis (PCA) of IS insertions across 2,124 *Y. pestis* genomes. **a,** Clustering of lineages based on the PCA of 2,566 chromosomal IS insertion sites identified within the analyzed genomes. Each point represents a genome, and ellipses mark areas including at least 95% of samples. Clusters of the three different 1.ANT clades are labelled. **b,** Focus on the clustering of different lineages from the PCA after excluding 1.ANT genomes. Labels mark the clustering of different lineages based on the chromosomal IS sites patterns on two Principal Components (PC). The upper panel shows PC1 vs PC2 and the bottom panel shows PC1 vs PC3, and percentages of variance explained by each PC are shown.

**Extended Data Table 1.**
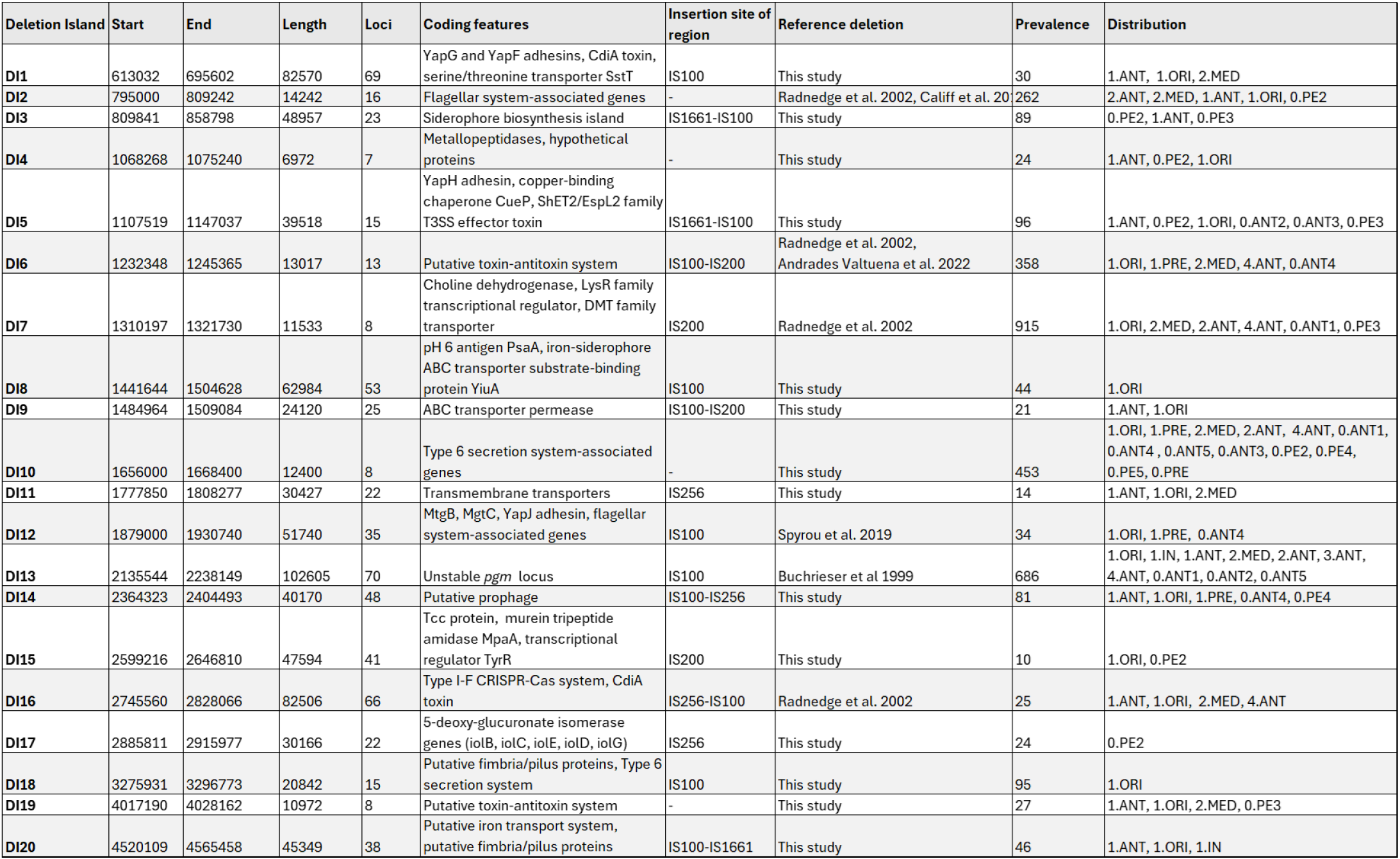
Features of the 20 identified deletion islands (DI) in the sequencing coverage analysis of 2,806 *Y. pestis* genomes. Approximated start and end coordinates of each large deletion are indicated in the positions within the CO92 reference chromosome.

