## Supplementary Information for "Global evolutionary patterns of *Yersinia pestis* and its spread into Africa"

### Supplementary Information 1

#### Historical contextualisation of plague branches' introduction and spread within Africa

##### The emergence and an early history of 1.ANT Branch: a historical hypothesis

As estimated in the present paper, the Most Recent Common Ancestor (MRCA) of all 1.ANT lineage emerged at some point between 1355 and 1528CE (median date: 1429), having diverged from Branch 1B. Historically, Branch 1B was responsible for what is known as the *pestis secunda* of 1356-66 – the second wave of the second plague pandemic, which radiated out of a putative reservoir in South-Central Germany and ravaged all over Europe, the Caucasus, the Middle East and North Africa[1–3]. Branch 1B itself was born as a result of the early split within the main Branch 1 (causing the Black Death), into two sub-lineages (Branch 1A and Branch 1B), seemingly in the course of plague's invasion of South-Central Germany in 1349, during the Black Death, or shortly hereafter[1].

During the *pestis secunda* wave, Branch 1B strains would eventually reach what can be regarded as geographic 'dead ends' – namely, regions constituting ecological or climatic hurdles to their further spread. By 1366, the wave seems to have reached the end of its quasi-global spread. Importantly, of over 100 sequenced genomes deriving from various post-Black Death contexts in Europe (chronologically extending from c.1357 to c.1771), only 9 genomes, all associated with different *pestis secunda* outbreaks, belong to 1B lineage, indicating that the branch appears to have migrated out of its putative reservoir in South-Central Germany – and Europe and possibly never returned there[1,3]. In these circumstances, Branch 1B strains could theoretically either die out at their geographic 'dead ends', or carry on evolving, having seeded new foci, whether in the course or at the end of its journey (at one of the geographic 'dead ends'). Given the subsequent evolutionary history of 1B and its offspring branches, including the oldest of them all - 1.ANT – it was clearly the latter, rather than the former scenario.

One of such geographic 'dead ends' was Egypt, where the *pestis secunda* is reported between February and July 1363. A systematic analysis of contemporary Mamluk-era chroniclers' description of subsequent plague waves indicates that albeit some of these waves were imported from outside (the Maghreb, Syria, or elsewhere in the Mediterranean), some are likely to have started in Egypt (possibly in Upper Egypt), before spreading to the north. Such waves are recorded in 1367-8, 1388-9, 1406-7, 1415-6, 1418-20, 1436-9, 1444, 1454, 1504-5 and 1513-4[4–6] – overlapping the estimated chronological timeframe of the emergence of 1.ANT MRCA, between 1355 and 1528CE (median date: 1429).

Hence, it is possible that there was a native Egyptian plague reservoir, from which the aforementioned waves were radiating, potentially associated with 1B lineage, and where the MRCA of the 1.ANT branch may have emerged, within the same reservoir. Although this hypothesis cannot be confirmed without ancient DNA (aDNA), it is partially supported by the argument, discussed in the next section, that Egypt was the most likely source of plague introduction into the Great Lakes region of East Africa, possibly in the late 17$^{th}$ century.

#### General Contours of Plague Evolution in Africa, c.1700-1920

##### 1. 1.ANT Branch in East Africa and beyond

*Plague's beginnings in Buganda c. 1700 and its possible source*

The only region in East Africa, which is known to have experienced plague outbreaks before the mid-19th century, is the Kingdom of Buganda, situated on the north-western shore of Lake Victoria. According to the oral tradition, collected and codified in the early 20th century, the earliest outbreaks in the kingdom go back to the reign of *kabaka* (king) Kayemba (c.1670-c.1700), and are associated with his son Kawumpuli, born limbless and expelled, because of his deformity, from his father's domains. After his death, Kawumpuli became the *lubaale* (national spirit) of Buganda associated with plague, with a dedicated temple in Buyego (near today's Bombo), and his name signifying 'plague' in Luganda language[7–9].

It is unclear, at this stage, how the plague was imported into Buganda c.1700 in the first place, with Ethiopia, the Swahil Coast, Sahara and the Nile being suggested as potential entry sources[9]. However, there is no historical evidence of any epidemics that could be associated with plague in the first three regions[10], in contrast with the Nile region, making it the most likely candidate of the four. Specifically, there was one environmental crisis event in the late 1690s that may have been a context, in which plague was introduced into Buganda.

To reach the Great Lakes, the pathogen had to traverse about 4,500km along the Nile from Lower Egypt, or some 3,700km from Upper Egypt, crossing all six cataracts and then delving into the difficult terrain of southern Sudan. Although, according to some early modern authors, plague was unknown to regions south of the second cataract[11,12], there was one exceptional case, when the pathogen appears to have crossed into Funj Sultanate (overlapping, roughly, with what today is Sudan) and potentially further south. The event in question consisted of an extreme drought of 1694-5, caused by abysmally low Nile levels (some of the very lowest on record) and evolving into a harsh famine, and an ensuing plague outbreak, attacking Egypt and Sudan in 1695-6[13–16]. Importantly, the same famine was characterised by an omnipresent human migration all over the Nile, with starving humans searching for food resources, and, with the plague outbreak, spreading the pathogen.

It was around the same time that the Lake Victoria witnessed a major migration of the Luo/Lwoo people from southern Sudan into what is now Nyanza Province of Kenya (on the north-eastern share of Victoria). Although the accurate chronology cannot be established, oral tradition had it that the same migration was triggered by drought, famine and invasions into Luo/Lwoo territories in the late 17th century[17]. The Luo/Lwoo migration would pass along the Nile, via the Shilluk, Dinka, Nuer and Anuak polities, and then into their new home on the Lake Victoria[18]. Taking this route would imply a contact and/or conflict with several Bantu- and Nilotic-speaking native peoples of the northern shores of the Lake Victoria – including the Iteso, who left their native territories in the Usuku district (northeastern Uganda) and migrated southwards into Bukedi on the north-eastern shores of the lake[17,19]. The westernmost edges of Bukedi are about 200km east the site of Kawumpuli's temple and about 125km east of the Nile's origin. Thus, the late 17th-century environmental crisis along the Nile, the migrations of the Luo/Lwoo and the Iteso, all appear to match the chronology of the earliest plague outbreaks in Buganda c.1700.

The 1695-6 plague outbreak in Egypt and Sudan falls within the upper limits of the Bayesian estimate of the emergence of the 1.ANT3 clade MRCA (1472-1697; median date: 1582). Given the subsequent phylogenetic history of 1.ANT branches in East Africa, it is possible that the 1695-6 outbreak, which may have been imported to Buganda, was associated with the emergence of the 1.ANT3 MRCA. However, with no aDNA from that region, there is no way to ascertain this hypothesis.

*Plague in Buganda: the endemic stage, c.1700-1850*

In the course of the 150 or so years since the beginning of Kawumpuli's cult c.1700, plague would persevere in Buganda, occasionally causing outbreaks in native rats and humans. Thus, there were plagues of rats during Kayabaggu's reign (c.1760-90), while under his successor *kabaka* Jungu (c.1790-1800), there were several plague outbreaks, leading to the desertion of many settlements. During King Semakokiro's reign (c.1800-12), there was an outbreak which killed, among other people, Kaduwamala, a deposed prime minister[8,20]. There is no evidence from oral histories of neighbouring regions that plague spread out of Buganda to other territories around the Great Lakes, and it is likely that it was confined to the former, persisting in a local reservoir in an endemic form.

This may be explained by the relative geographic socio-political isolation and economic autarky of different Interlocustine polities. In the case of Buganda, it was not until Junju's reign (1790-1800) that we witness the first signs of nascent inter-regional trade with the kingdom of Karagwe (north-western Tanzania), based on cotton and cowrie shells in exchange for Buganda ivory[20]. These commercial relationships continued under Semakokiro (c.1800-12)[7,21,22], but it was not until the arrival and establishment of Arab merchants from Khartoum and Oman in the Great Lakes region in the 1840s-1850s that the geographic range of trade expanded considerably, and more and more regions were gradually integrating into medium-distance trade (see below). Likewise, there is no evidence of any large-scale human migration, caused by either conflict or subsistence crisis, in the region in the pre-1850 period.

The geographic restriction of plague to Buganda in the 18th and the first half of the 19th century may be reflected in the phylogenetic reconstruction of the early evolutionary stages of Branch 1.ANT history. The environmental crisis of 1694-6 and the initial importation of plague into Buganda c.1700, could be the very context in which 1.ANT3 may have emerged – possibly in a newly established reservoir not far from Kawumpuli's temple. Between the emergence of 1.ANT3, whose strains have been documented only in eastern DRC and Kenya (with Buganda situated exactly halfway between the two), and the diversification of 1.ANT sub-branches (between 1711 and 1882; median date: 1802), the same branch appears to be characterised by a genetic monophyly, which, in turn, may reflect its geographic restrictiveness to Buganda.

*Plague in East Africa: the expansive stage, c.1850-1900*

The second half of the $19^{th}$ century was, by contrast, characterised by the geographic expansion of plague in the Great Lakes region. In the 1850s, plague penetrated, for the first time, into Busoga, Ankole, Buddu, Karagwe and Kisumu regions. Around 1875, plague outbreaks occurred in Ishanze (Nkore/Ankole region in Uganda) and Kanyigo (northern Kiziba in northern Tanzania) (Lwamgira 2020: 324-5). In 1886, plague appeared in Imagi and Uhehe (Tanzania), having been introduced from the north during Wahehe chief Mkwawa's campaigns[23]. From Uhehe, the pathogen expanded southwards to Ubena and Ilembula (South-West Tanzania, close to the Malawi border), where it broke out in 1891 and 1893[23,24], while in 1900, the disease reached the Mpwapwa and Dodoma regions in Central Tanzania[24].

At the same time, the disease was making itself known in the north, throughout what is today Kenya. In 1892-3, there was a plague outbreak around Ndara Hill and Sagalla[25]. The disease then spread into Machakos (some 60km southeast of Nairobi) either in 1895 or 1897, before carrying on to the Taita-Taveta region in 1897-8 (some 190 km north-west of Mombasa)[26,27]. In 1898, the plague finally reached the Swahili Coast, where a plague-infected passenger was identified in Mombasa, on the board of a German ship bound for Zanzibar[26]. Concurrently, the plague was also spreading in the west: between c.1888 and c.1898, it crossed Lake Albert into the Nizi region of eastern Congo[28].

The geographic expansion of plague in East Africa c.1850-1900 was undoubtedly facilitated by the commercialisation of the region, first with the arrival of Arab merchants of Khartoum and Oman, and later of Gujarati traders, as well as European missionaries, merchants, explorers and settlers[29,30]. Initially, the two main export items were ivory and slaves, although the demand for the latter declined a great deal with anti-slavery legislation of colonial administration in the 1870s-80s. In exchange, East African natives would receive Gujarati cotton, European firearms and a handful of other goods[31,32]. In absence of pack-animals, a central role in transportation and communication was filled by human porters, who were considered not only the carriers of merchandise, but also of plague[33–35].

The commercialisation of the Great Lakes region and its integration into the wider Indian Ocean and Nile trade systems was not the only factor contributing to the increase in human migration. Further, it was facilitated by the creation of agricultural plantations by colonial authorities, both British and German, and the construction of railway systems: German Usambara Railway, connecting Tanga in the east with Moshi in the west (running 350km), and British Uganda Railway, running from Mombasa in the east to Kisumu in the west (1,060km)[36,37].

Furthermore, we have to account for military conflicts, whose volume rose a great deal in the second half of the $19^{th}$ century, as another channel of plague spread. Thus, c. 1876, King Mutesa's soldiers raided Bukedi region (the north-eastern shore of the Lake Victoria), only to get infected with plague and bring it back into Buganda[38]. In 1880, another plague outbreak occurred in Mutesa's solders during their campaign in neighbouring Busoga[22]. Similarly, the Uhehe region of Central Tanzania experienced the first plague outbreak in 1886, in the context of Chief Mkwawa's invasion there[23].

The diversification of 1.ANT lineages would occur precisely in this economic and socio-political context. The first branch to diversify was 1.ANT1 (between 1711 and 1882; median date: 1802), most likely in the context of the early plague spread around the northern shores of the Lake Victoria – which is corroborated by the phylogeography of its strains, known to be present only in East DRC and Uganda. This was followed by the respective diversification of 1.ANT2 (between 1772 and 1920; median date: 1851) and 1.ANT3 (between 1800 and 1920; median date: 1873), in the context of the piecemeal expansion of *Y. pestis* eastwards and southwards. Indeed, while 1.ANT2 strains are found in Tanzania, Kenya and Yemen, the 1.ANT3 ones are attested in East DRC and Kenya.

*Plague in the Indian Ocean and southern Africa: the expansive stage, c.1900*

Following its presence in Mombasa 1898, the plague invaded Oman in 1899, and Yemen in the following year[39,40]. Once in Yemen, the plague may have seeded a reservoir in the Asir mountains (on the Yemenite-Saudi border), associated with the origins of the 1969 outbreak[41], whose genome has been sequenced in this study and belongs to 1.ANT2 clade (genome id IP1865H). Somewhat later, the pathogen began moving southwards, spreading into Malawi and the Eastern Province of Zambia in, respectively, late 1916 and early 1917[42].

#### 2. The Introduction of 1.ORI5 Branch into North and West Africa, 1898-1912

There is, however, no evidence that 1.ANT strains were migrating further south: all the genomes from South Africa, Namibia and Zimbabwe sequenced in the present study are positioned on 1.ORI5 branch. The same branch is associated with the third plague pandemic commencing in Hong Kong in 1894 and spreading into Mozambique, Transvaal in 1899, via Mauritius and Madagascar (where it had been attested in 1898), and into Cape Town a year later[43,44]. It was not, however, until the 1930s that the disease would spread further inland – first into Namibia (1931) and then Lesotho (1935); no human plague has been reported in Zimbabwe until 1972[24,45].

Concurrently, plague was re-introduced into North Africa, with the first cases reported in Alexandria and Philippeville/Skikda (Algeria) in early May 1899. As far as Alexandria is concerned, the disease appears to have been introduced on pilgrimage ships from Jeddah, where a harsh outbreak occurred in conjunction with the annual *hajj* [46,47]. The provenance of the Philippeville's outbreak is unclear, but either Jeddah or a west Mediterranean port (Lisbon or Porto) could be a potential candidate[48,49]. Importantly, plague had initially been introduced into Jeddah by Indian pilgrims two years earlier (1897) from Mumbai/Bombay, where, in turn, it had been introduced in 1896 from Hong Kong, in the context of the commencement of the third plague pandemic[50]. Given the association of the Hong Kong outbreak and the ensuing wave with 1.ORI5 strains, we may infer that the Mumbai/Bombay, Jeddah, Alexandria and Philippeville outbreaks were, too, caused by strains of the same branch.

In 1907, the plague reached the port of Tunis, most likely from Marseille[49]; the Marseille outbreak, too, was associated with the third plague pandemic wave[51], and hence, likely with Branch 1.ORI5.

On 20 July 1909, plague was reported in the Chaouia region of Morocco. Unlike the Philippeville and Tunis outbreaks, the disease seems to have been introduced by the Bedouins from the Tafilalt region, some 420km south of Casablanca[52,53]. Given the putative northbound movement of the 1909 outbreak in Morocco (from the Tafilalt region into Casablanca, then Rabat and Tangier in the course of 1910-1), we may presume that 1.ORI5 strains got focalised somewhere in the Sahara at some point between their initial introduction in 1899 and the Moroccan outbreak of 1909. In 1912, plague reached the port of Ziguinchor in southern Senegal, most likely on a ship from Casablanca; in April 1914, the epidemic broke out in Dakkar[49,54].

Ever since its (re-)introduction into West Africa, plague became an endemic disease in that part of the continent, having established multiple foci in different regions and causing outbreaks in humans incessantly until the mid-20$^{th}$ century (Algeria: 1899-1900, 1902-4, 1907, 1911-3, 1916-31, 1935-7, 1939-40, 1944-6, 1950; Tunisia: 1907, 1910, 1920-31, 1933-41, 1944-9; Morocco: 1909-3, 1919, 1921-4, 1929-35, 1940-5 and 1948; Senegal: 1912, 1914-5, 1917-45)[24]. After some half century of respite, an Algerian focus got reactivated in 2003[55].

##### 3. The Introduction of 2.MED1 Branch into Libya (c.1913?)

Three genomes from Tobruk (North-Central Libya) associated with a 2009 outbreak are all positioned on 2.MED1 branch. Up until now, strains associated with the branch in question have been attested over a very wide geographic range - from northern Xinjiang and western Mongolia in the east to the Caucasus in the west, via lowland steppe and semi-desert reservoirs of South Russia, Kazakhstan, Turkmenistan and Uzbekistan and highland zones of eastern Caucasus, the Kurdistan province of Iran, Tian Shan and Altay. These 2.MED1 reservoirs are being sustained by a wide variety of wild rodents, including common voles, jirds, gerbils, marmots and susliks. In this regard, the Libyan genomes are geographic outliers, and their origins are to be sought somewhere to the North-East of Libya.

The present study has estimated the emergence of the Libyan clade at some point between 1911 and 1947 (median date: 1937), placing it firmly in the earlier 20$^{th}$ century. Importantly, after the outbreaks in 1858-9 and 1874, there is no evidence of human plague in Libya until its re-emergence in 1913, leading to the 1913-22 wave and subsequent outbreaks in 1924-34[24,56,57]. Hereafter, we hear of no human plague again until 1972, followed by the outbreaks of 1976-7, 1984 and 2009[24,58]. Hence, we may fairly safely associate the emergence of the Libyan clade of 2.MED1 with the period of 1913-34. There are two possibilities: (1) the clade emerged in association with the 1913-22 outbreak, either if introduced from elsewhere or if spilling over from a local endemic reservoir, if there was such; (2) the clade rose after the 1913-22 wave, if the strains introduced in 1913 were associated with a more ancestral clade, found in Central Asia and Caucasus.

Regardless of the scenario, it is essential to establish the proximate origins of the 1913-22 wave, commencing shortly after the Italian takeover of Libya from the Ottomans in 1911. The 1913-22 wave commenced with outbreaks in the ports of Tripoli and Derna in, respectively, 1 and 15 July 1913. In Benghazi, the plague was reported first in December of the same year lasting from 1 July until 30 September 1913[59]. In Janzur, the plague was reported in July 1914; concurrently, the disease spread westwards into Tunisia[60]. During the same wave of 1913-22, Libyan ports witnessed recurrent plague introductions causing widespread outbreaks, with 1917 being a particularly harsh year.

Importantly, plague appears to have seeded a local reservoir, in one of the desert regions south of Benghazi, and it was from that reservoir that the 1858-9 and 1874 outbreaks most likely originated[56,57]. Importantly, the same outbreaks were confined to the Benghazi region only; by contrast, the 1913-22 wave commenced in Tripoli, being imported from outside, implying that it was associated with a different origin and a different reservoir. The present study has estimated the emergence of the MRCA of the Middle Eastern-Libyan 2.MED1 sub-clade to c.1749-1865 (median date: 1812), implying that the clade in question had emerged somewhere in the Middle East some 100 years before its introduction into Libya in the early 20th century. To reach Tripoli from the East, the plague would need to travel from Egypt (from Alexandria, to be specific), where, in turn, it would have to enter either by sea from the Persian Gulf and the Red Sea, or by land, via East Anatolia and Syria. However, with the exception of a very small-scale outbreak in Beirut in 1909-10, there is no evidence of any plague outbreak elsewhere in Syria in 1913 or preceding years[61]. By contrast, there were annual plague outbreaks in Egypt, with no respite, since 1899[62]. On the eve of the plague's introduction to Tripoli in July 1913, plague was present in Alexandria since 28 May (until 28 October) 1913[59] and it is most likely that it was from there that it entered Libya.

As noted, the initial reintroduction of plague into Egypt in 1899 was associated with the same wave commencing in Hong Kong in 1894, reaching India in 1896 and Jeddah in 1897, from where it reached Alexandria two years later[62]. Although there is, at present, no DNA associated with the 1899 introduction, one may infer that the outbreak in question was caused by 1.ORI5, rather than 2.MED1 strains. Following the 1899 introduction, plague appears to have become endemic to Egypt for the next 48 years[62].; however, in addition to local reservoir/s, there were recurrent introductions from outside, via the Suez Canal, or directly into Port Said or Alexandria[63,64]. Given the presumed eastern origins of 2.MED1, and the lack of plague activity in Anatolia and Syria, we would, as noted above, expect its strains to be introduced into one of Egyptian ports from somewhere in the Middle East, by the way of the Persian Gulf and the Red Sea.

One of the frequently shipping routes was that connecting Basrah and Bushehr, both on the northern limits of the Persian Gulf, with Port Said and Alexandria. Both Basrah and Bushehr were important trading partners of Egypt, shipping annually large volumes of goods to and from local ports[65]. Between 1910 and 1913, there were several plague outbreaks in Bushehr. Of these, the 1910 and 1912 ones appear to have been imported from Basrah[61,66] (which, in turn, would seemingly receive these from Jeddah, in the context of Hajj pilgrimages)[61,64,66] while the 1913 one, occurring in April-May, appears to have originated in Kurdistan earlier that year[67]. It is possible, then, that when plague reappeared in Alexandria in late May of the same year, it was reintroduced from Bushehr – and that the same strains originating in Kurdistan earlier that year were introduced into Tripoli a month later and seeded a new reservoir in Libya hereafter.

The Kurdish origin of the Libyan clade of 2.MED1 is corroborated by the phylogeography of the same branch, and the geographic trajectory of plague outbreaks in 19th- and 20th-century Iran. Although, as noted above, it has established itself over vast regions of Asia – from South Russia to Mongolia, via the Middle East and Central Asia -, this branch is notably widespread in Kurdistan. The existence of a plague reservoir in Iranian Kurdistan has been mapped and studied since the late 1940s by Marcel Baltazard and his colleagues[68], but noted already in 1907 by Ukrainian plague scientist Danylo Zabolotnyi, who was the first to note that plague waves witnessed in Mesopotamia and Iran tended to commence in Kurdistan[69]. For the purpose of the present study, by ‘Kurdistan’ it is meant a region overlapping with historically Armenian and Kurdish regions of South-East Turkey, and Kurdish regions of North Iran and North-East Iraq. However, another possibility is that the reservoir in question was situated somewhere in a region overlapping with today’s two Azerbaijan provinces of Iran, situated to the north of Kurdistan. Grossly oversimplifying, and in absence of a precise location of the reservoir(s) in question, we may refer to it as a ‘North-Western Zagros reservoir’. Indeed, the trajectory of plague spread can be established for the waves commencing in that putative reservoir(s) in 1771, 1798, 1829, 1833, and spreading into different regions of Anatolia, Mesopotamia, Iran and Caucasus[70–75]. The waves commencing in 1871 and 1876 were limited to parts of Iran, Iraq and Caucasus[72–74,76]; the 1913 outbreak spread only in eastern Iran[59,67]; 1947, 1951-2, 1961, 1963 and 1966 outbreaks were confined to Iranian Kurdistan[77], while the 1958 one to West Azerbaijan province of Iran[77].

### Supplementary Information 2

#### Additional supplementary figures

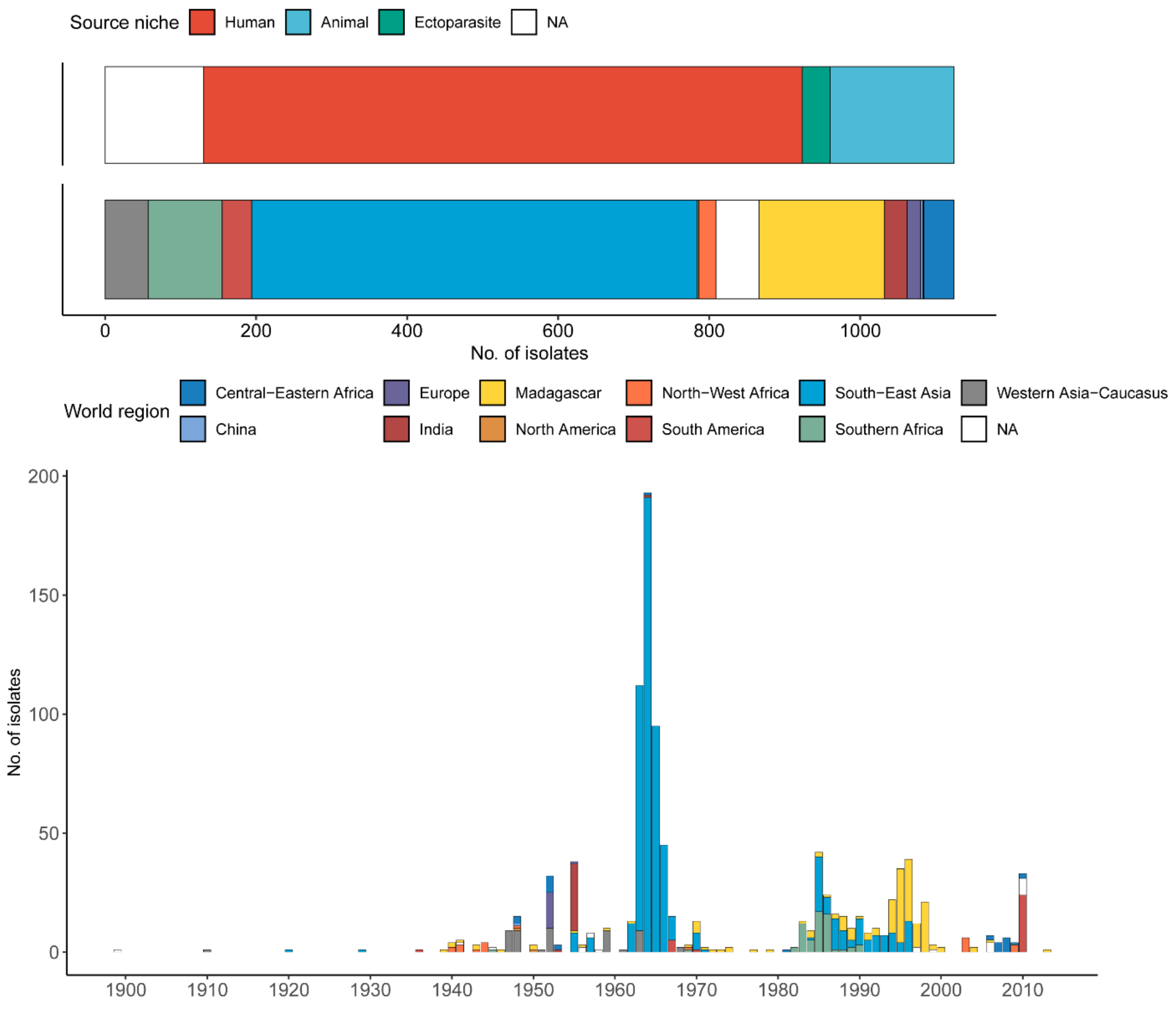

Supplementary Fig. 1 - Distribution of *Y. pestis* isolates sequenced in this study, by source type, geographic origin and isolation year (*n*=1,124). Plots were generated using the ggplot2[78] package in R 4.4.0[79].

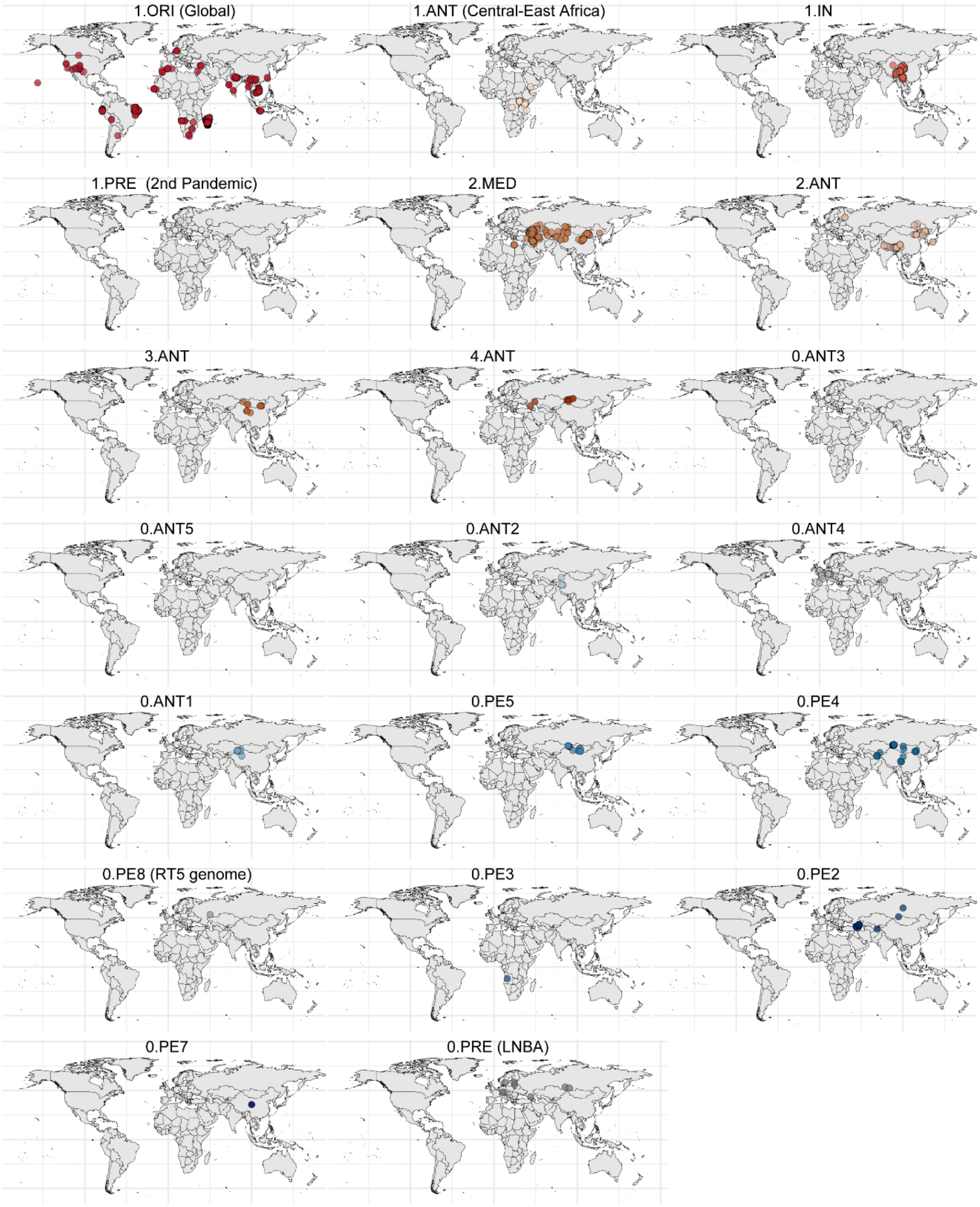

Supplementary Fig. 2 - Geographic distribution of *Y. pestis* genomes for each of the 20 main phylogenetic lineages of the species. Precision up to town level were used for the coordinates of each genome when available, while centroid coordinates were used for samples for which only the country of origin was known.

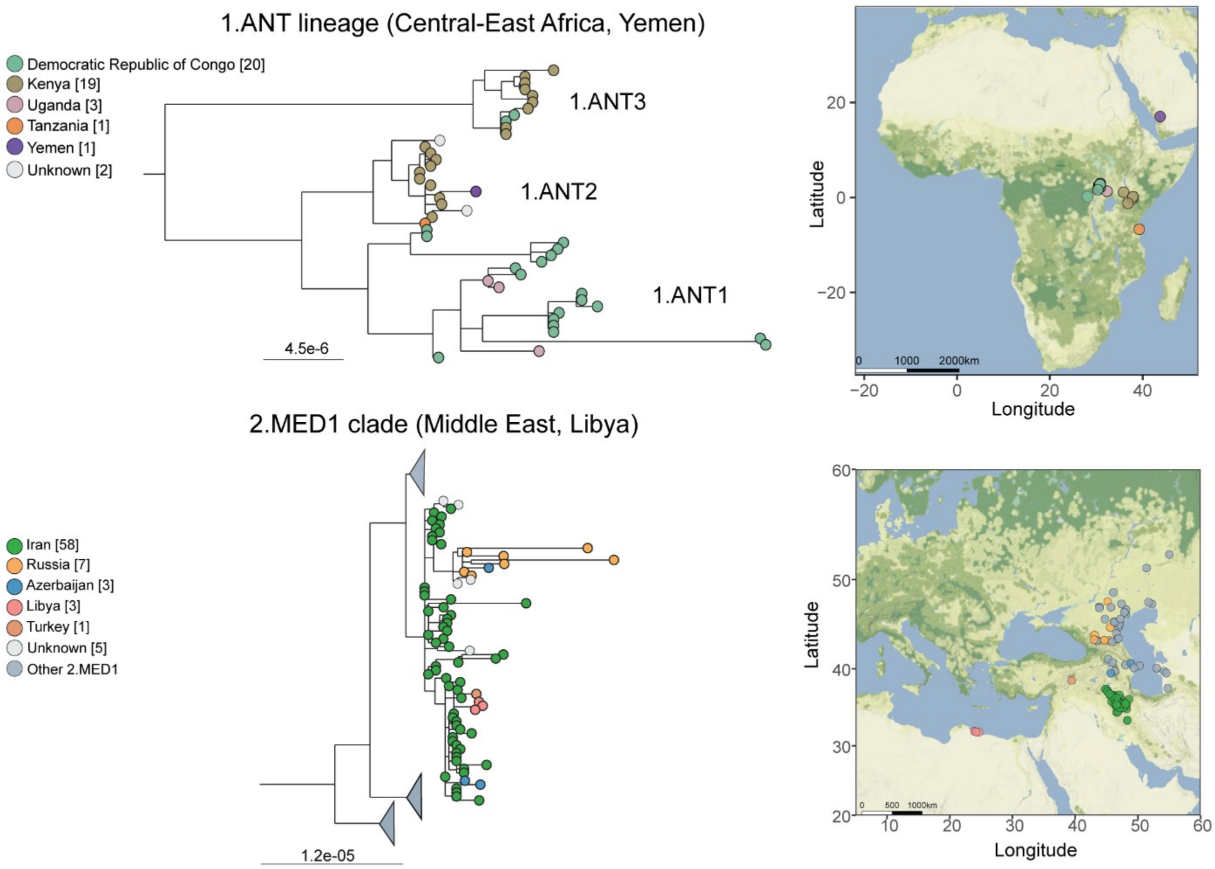

Supplementary Fig. 3 - Phylogeny and geographic distribution of 1.ANT and 2.MED1 clades. Upper panel: Subtree corresponding to the 1.ANT lineage. Lower plot: Subtree corresponding to the 2.MED1 clade, focusing on the subclade that includes isolates from Western Asia (Kurdistan, Iran and Turkey), Caucasus (Russia, Azerbaijan) and Libya, while the other 2.MED1 subclades are collapsed. Both panels have been extracted from the complete species maximum-likelihood phylogeny displayed in main Fig. 1, inferred from RAxML-NG 1.0.1[80]. Maps were generated using ggmap R-package[81]. Tips and dots on the maps are coloured according to the country of origin of the bacterial isolates. Scales under each phylogenetic tree represent the number of substitutions per genomic site.

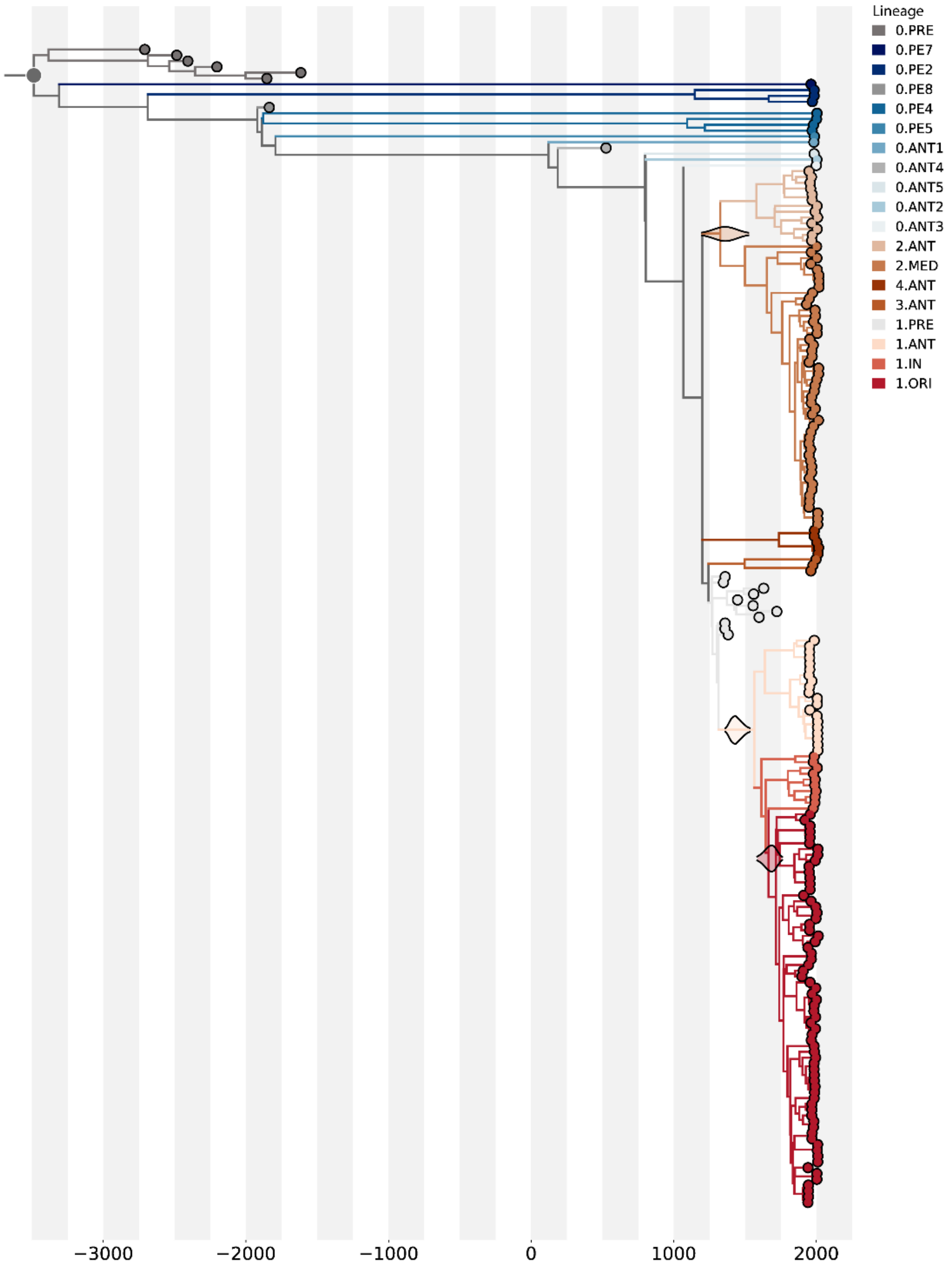

Supplementary Fig. 4 – Time-scaled Maximum Clade Credibility (MCC) tree from the molecular dating inference using BEAST v2.6.6[82]. The dating dataset includes 200 modern and ancient *Y. pestis* genomes, and *Y. pseudotuberculosis* IP32953 genome to root the tree (branch not shown to facilitate visualization of the divergence dates within the *Y. pestis* lineage). The tree is scaled to inferred median divergence dates, from the most recently sampled isolate (2018) to the most recent common ancestor (MRCA) of most ancient *Y. pestis* (0.PRE) and the rest of lineages (median year: -3,486, 95% posterior interval year: -2,927; -4,203). Labels under the axis represent the divergence year. The 95% highest posterior density (HPD) distributions for the date of the MRCA of 2.MED, 1.ANT and 1.ORI lineages are represented with violin plots centred on each MRCA node. The tree figure was visualized in Python3 using the Baltic 0.3.0 library[83].

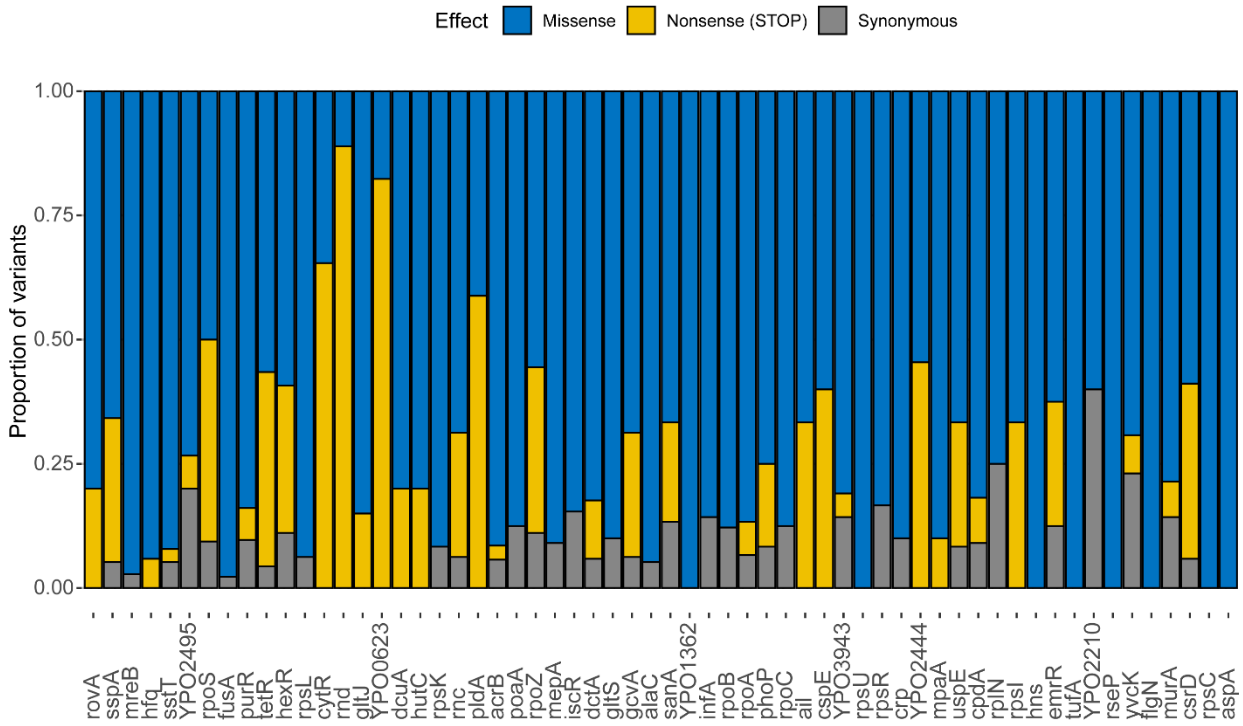

Supplementary Fig. 5 – Effect of the variants within the 60 genes presenting a number of substitutions significantly deviating from the theoretic expectations under neutral evolution, identified from the genome analysis of 2,700 *Y. pestis* isolates. Barplots represent the proportion of mutations that cause amino acid substitutions (missense), premature stop codons (nonsense) or without amino acid change (synonymous).

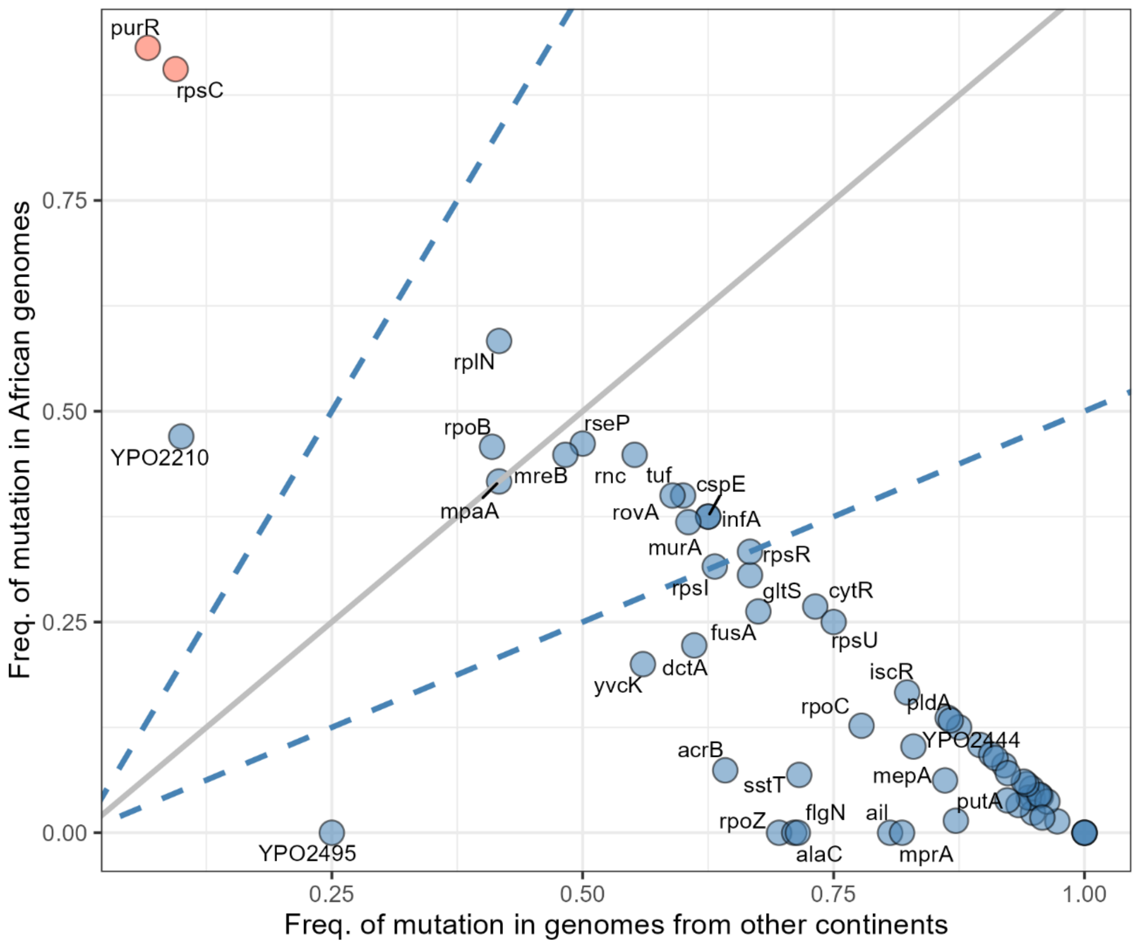

Supplementary Fig. 6 - Allele frequencies in 60 identified genes presenting significantly higher mutation density. The scatterplot represents the proportion of genomes from Africa (y axis) and other regions (x axis) presenting at least one mutation within the gene, from the total genomes presenting at least a mutation within the same gene. The dotted blue lines indicate where the allele frequencies in African genomes are two-fold higher than in other continents (upper line) and where the frequencies in genomes from other continents are two-fold higher than in African genomes (lower line), while the solid grey line indicates when African and other continents allele frequencies are equal. Dots coloured in red highlight genes with mutations emerging mainly (>90%) in African genomes, while presenting low mutation frequency (<10%) in genomes from other regions.

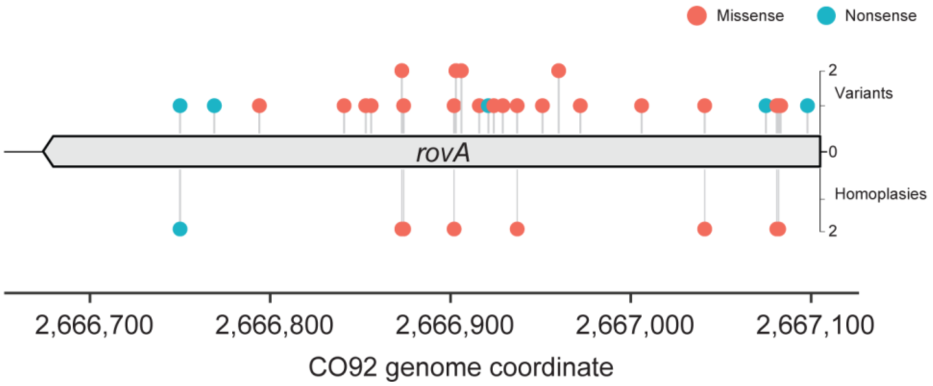

Supplementary Fig. 7 - Gene map highlighting the frequency of missense and nonsense mutations, and the frequency of homoplastic mutation events identified within *rovA* locus from the analysis of 2,700 *Y. pestis* modern genomes. Source data is provided in Supplementary Table 9. The scale represents the chromosome coordinates corresponding the *Y. pestis* CO92 reference (RefSeq ID: NC_003143).

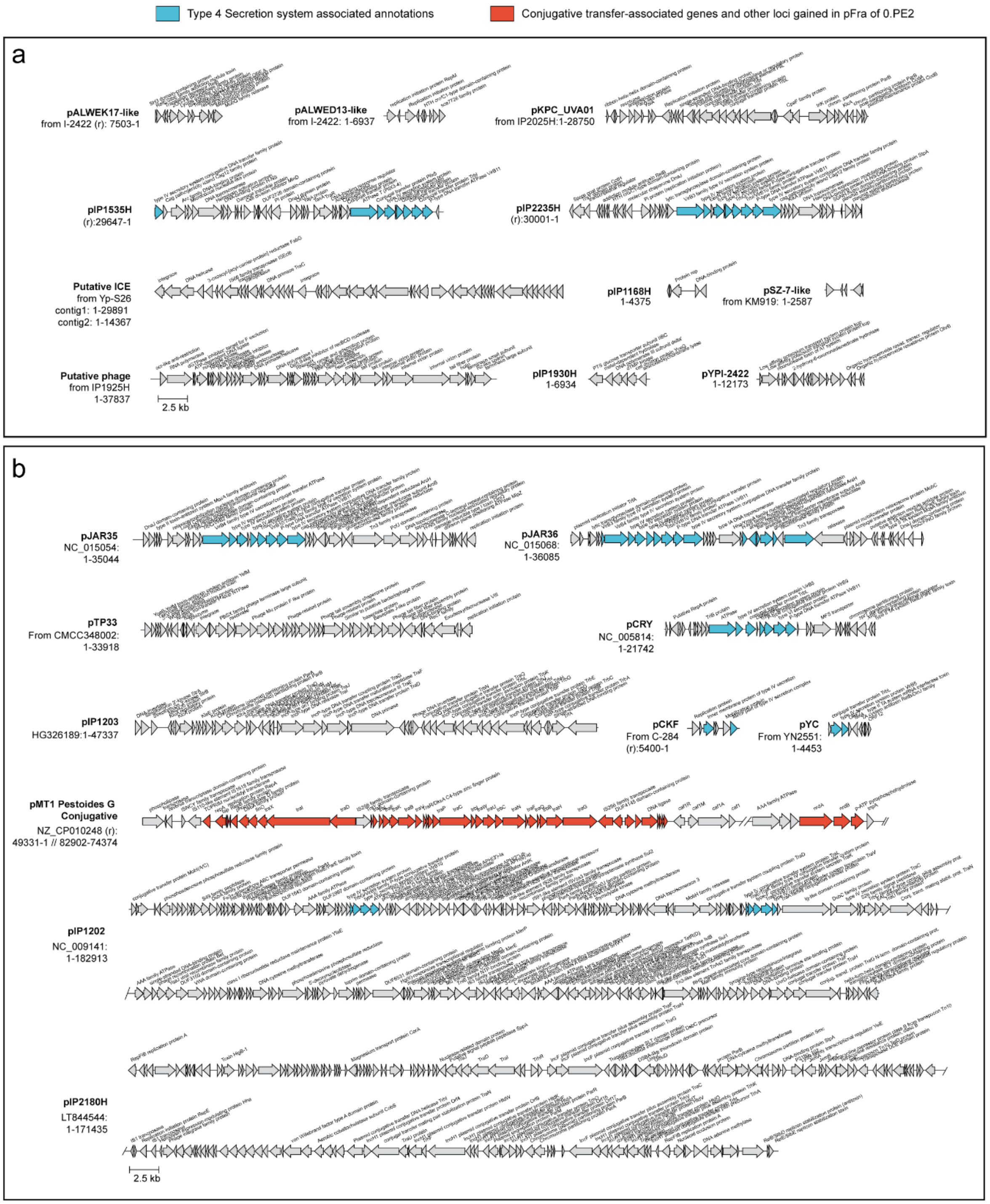

Supplementary Fig. 8 – Gene maps of the total 21 likely acquired genes identified in modern *Y. pestis* assemblies. a, Schematic representation of novel mobile genetic elements identified in this work, including 9 plasmids, a putative integrative and conjugative element (ICE) and a putative phage. b, other replicons and likely acquired gene clusters in modern assemblies previously described in the literature. Assembly identifier, when available, is indicated. Visualization of annotated clusters of genes was performed using Clinker 0.0.30[84].

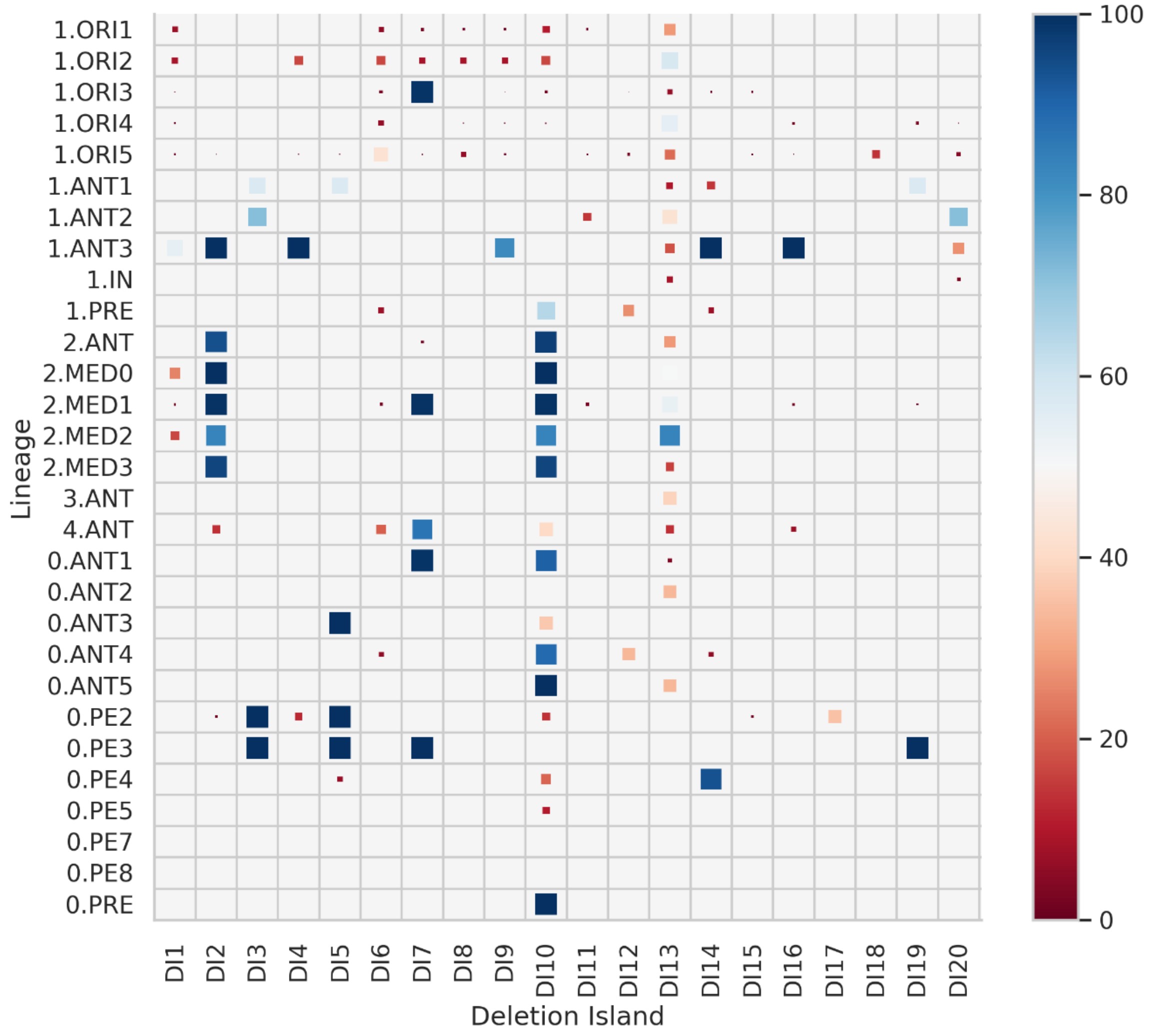

Supplementary Fig. 9 - Distribution of 20 deletion islands (DI) across *Y. pestis* clades. The heatmap shows the prevalence of large deletions within each clade based on the analysis of 2,806 genomes, with square sizes and colour scale adjusted to their value in percentage.

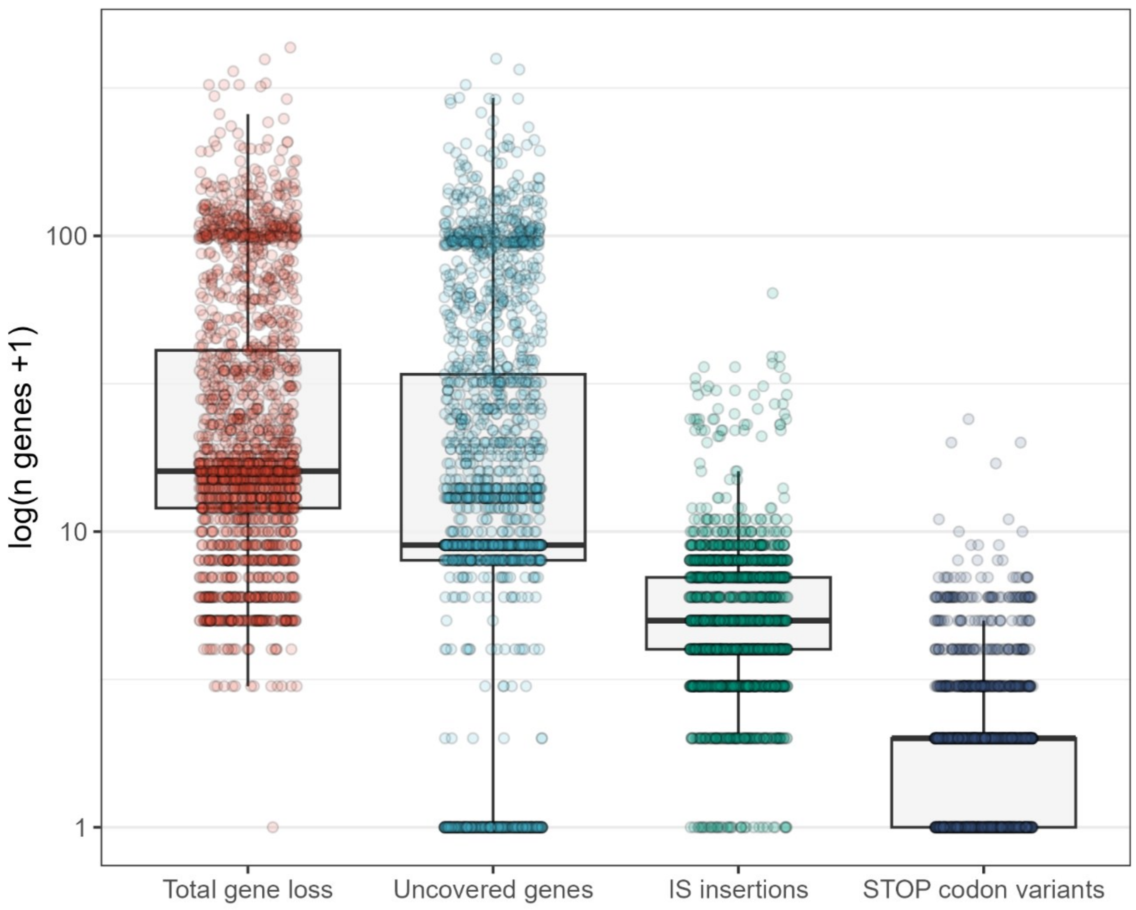

Supplementary Fig. 10 - Mechanisms driving gene loss in 2,128 *Y. pestis* genomes. Boxplots showing the distribution of the total number of genes lost or inactivated per genome (left distribution) and classified according to the inactivation mechanism. Uncovered genes represent genes without sequencing coverage; IS insertion represent genes in which the insertion of an IS element was inferred for the genome, causing their pseudogenization; STOP codon variants represent genes inactivated through the emergence of at least one nonsense mutation causing the emergence of a premature STOP codon. The boxplots represent the 75$^{th}$ percentile, the median (bold line) and the 25$^{th}$ percentile for the distribution of total genes lost per genome, while values beyond 1.5 times the interquartile range are considered outliers.

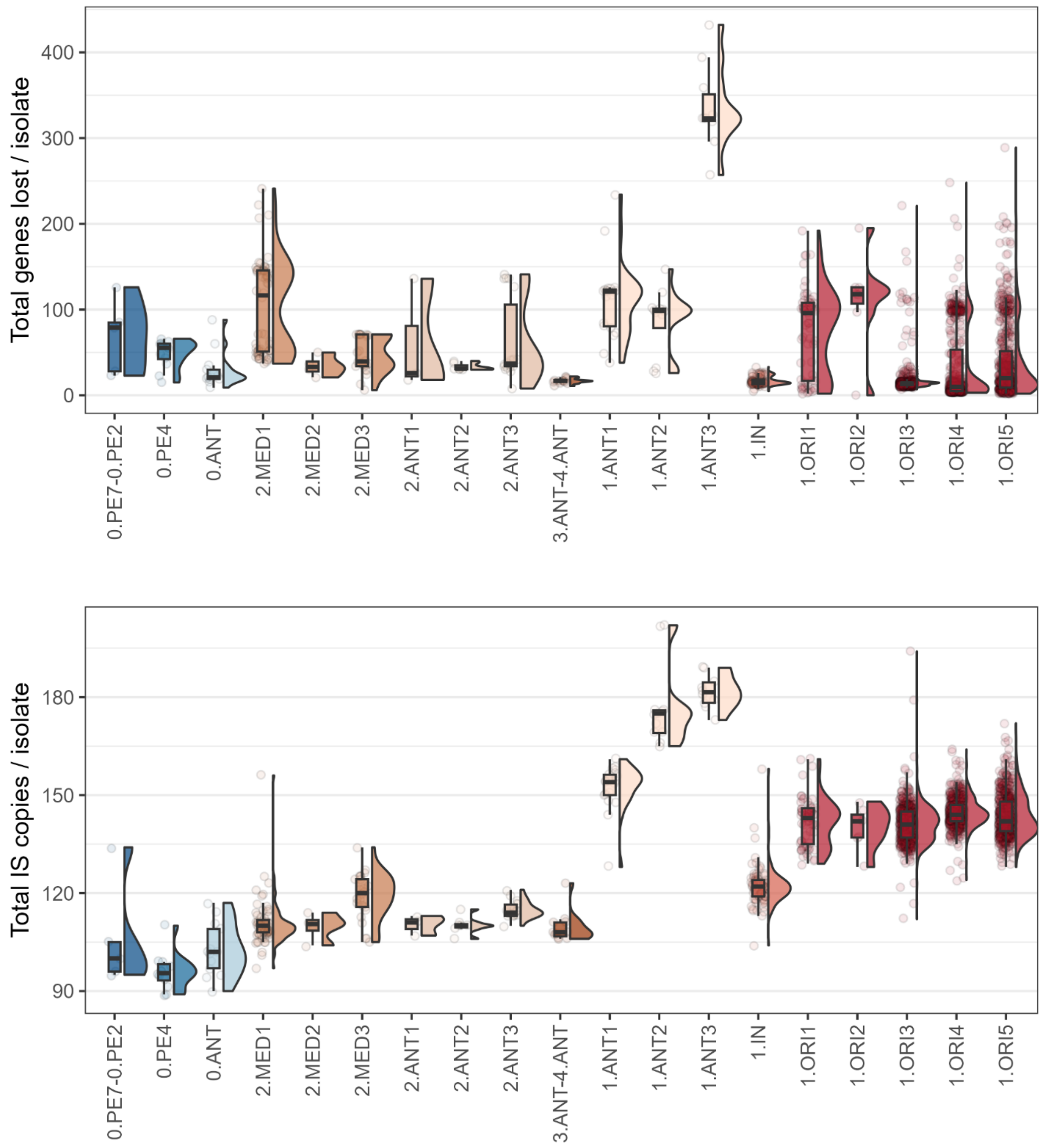

Supplementary Fig. 11 - Gene loss and IS dynamics in *Y. pestis* evolution. Distribution of total counts of lost genes (upper plot) and IS copies (lower plot) per genome across different phylogenetic clades. The box plots mark the 75$^{th}$ percentile, the median (bold line) and the 25$^{th}$ percentile for the distribution of total genes lost and IS counts per genome in each plot, respectively, while values beyond 1.5 times the interquartile range are considered outliers.

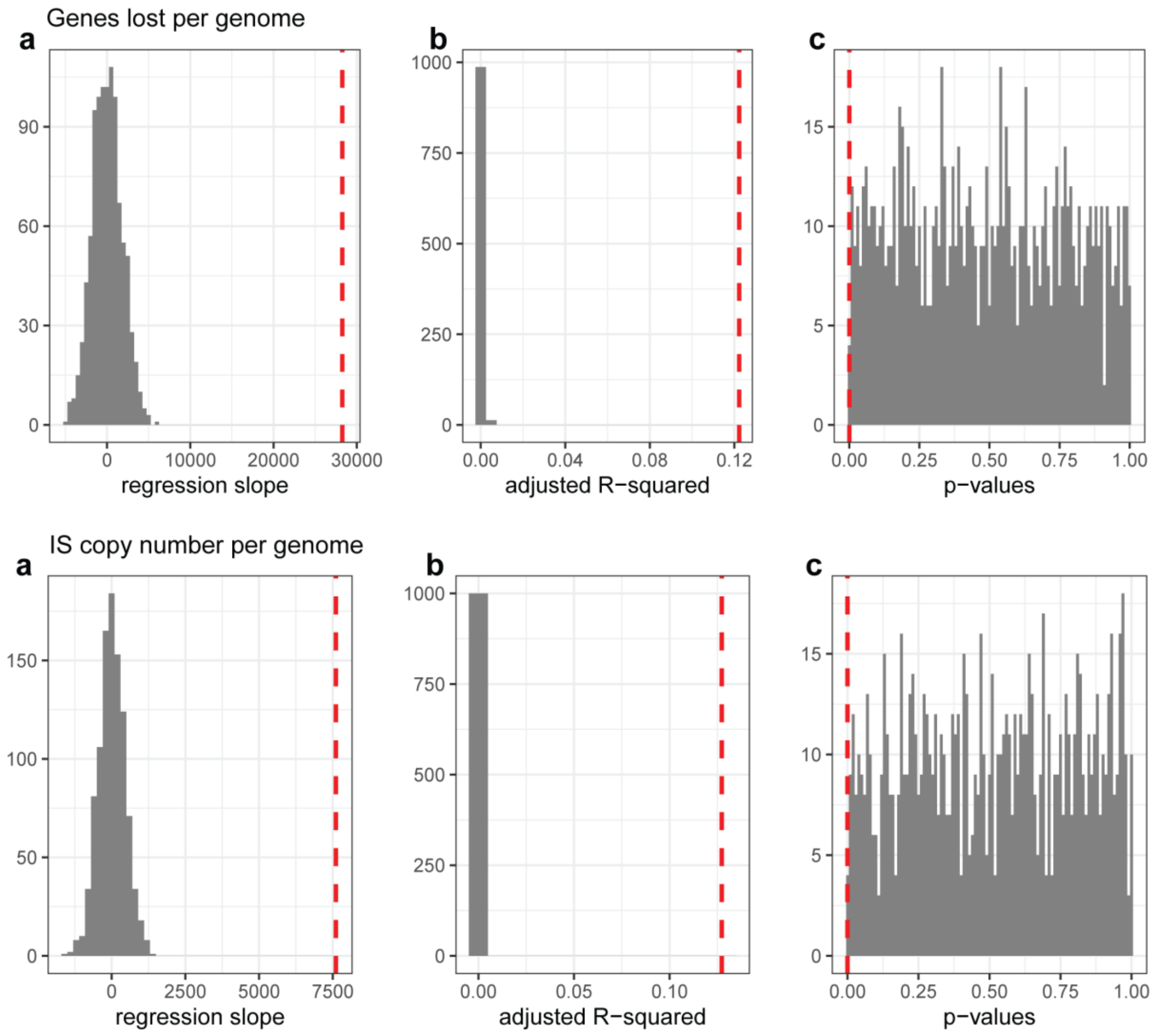

Supplementary Fig. 12 – Permutation tests of the linear regression of total genes lost (upper panel) and IS copy number (lower panel) vs patristic genetic distances in *Y. pestis* genomes. **a** (on both panels), regression slope values based on 1000 permutations (grey) with the real slope indicated by the red line. **b** (on both panels), same as panel **a**, but displaying the adjusted R-squared values for the 1000 permutations. **c** (on both panels), as in panel **a**, but showing p-values for the 1000 permutations.

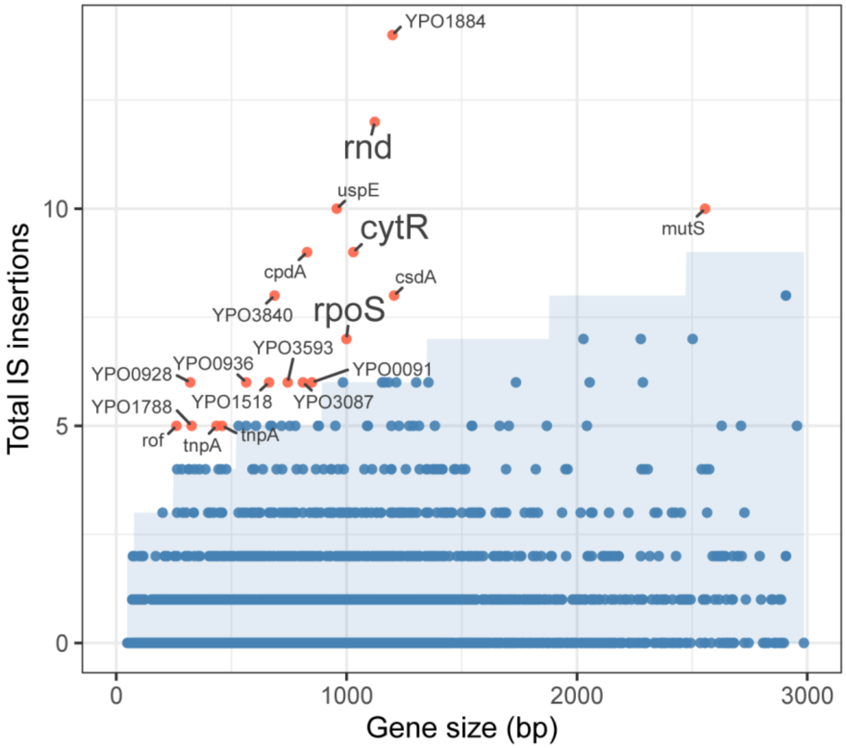

Supplementary Fig. 13 – Identification of IS insertions within genes. Scatterplot of IS counts within each gene sequence against the length of each locus in base-pairs. Coordinates for the chromosomal IS insertions were identified in 2,127 *Y. pestis* genomes using ISMapper 2.0.2[85]. A Bonferroni-corrected Poisson test was used to compare the observed number of IS insertion sites with the expected value for each gene given their sequence length. Dots in red highlight 19 loci with observed IS insertions sites significantly higher than expected under the Poisson distribution. The shaded area represents the expected number of IS insertions per gene under the assumption that IS insertions follow a uniform rate across the genome, given a starting significance level of 0.05. Highlighted labels indicate three genes also displaying increased rates of nonsense variants identified in the mutation analysis.

### Supplementary Information 3

#### Methods

##### Identification of genes with mutation counts deviating from theoretical expectations

The accumulation of nonsynonymous mutations has been associated with diversifying selection acting on the evolution of some genes[86]. To identify genes presenting a significantly higher mutation density in *Y. pestis* evolution, we adapted the procedure proposed by Cui et al., which compares the distribution of mutations per gene to theoretical expectations[87]. First, high-quality calls were obtained using RedDog[88] pipeline with default parameters from the mapping of 2,700 genomes representing all modern lineages of the species on the CO92 reference chromosome, and only variable positions covered in at least 90% of samples were kept. A consensus sequence integrating the identified variants was then generated for each genome. Sites potentially associated with intraspecies recombination were inferred from the whole-genome consensus sequences using Gubbins 3.3.5[89] and filtered out from the variant table (n=4,017 mutations). Genes in which at least 50% of their mutations are observed in only a single genome were identified and excluded (n=57 genes), as they may represent artifacts (i.e.: sequencing or assembly errors) or a diversity exclusive of a single sample, which does not reflect evolutionary pressures on the locus acting at the population level. After these filtering steps, the final dataset contained 2,546 genes. In total, 15,865 mutations were kept, from which 8,895 are nonsynonymous, 2,911 synonymous, and 4,059 are found in regions other than protein-coding ones. The total counts of each mutation category were summarized for each gene using the script countMEbyGene.py. We focused only on nonsynonymous mutations (n = 8,895), as they mainly represent changes in protein function. Gene lengths were extracted from the reference CO92 chromosome (NC_003143.1).

We then applied a $\chi^2$ goodness-of-fit test to compare the observed nonsynonymous mutation counts in each gene with a theoretical neutral expectation, following the methodology of Cui et al.[87]. We first computed the expected mutation rate per nucleotide as $\mathrm{m}=\mathrm{n}/\mathrm{N}$, where $n$ is the total number of nonsynonymous mutations and $N$ is the sum of all gene lengths. For each gene, the expected number of mutations was calculated as $mL_i$, where $L_i$ is the gene length. The test statistic was then computed as:

$$\chi^2 = \sum_{i=1}^{K} \frac{(n_i - mL_i)^2}{mL_i},$$

where $K$ represents the number of genes. The $\chi^2$ statistic was then computed by comparing observed and expected values across all genes. We estimated a large statistic ($\chi \approx 9668.62$), resulting in an extremely small *p*-value (of the order of $10^{-778}$), indicating a significant deviation from neutral expectations. Given that some genes may have expected mutation counts lower than 5 (a requirement for the $\chi^2$ test), we also drew a non-parametric distribution of mutation counts per gene as comparison. We simulated the distribution of mutation counts using a binomial model, where for each gene, we sampled 100,000 values from a binomial distribution with parameters $L_i$ and $m$. This allowed us to generate a non-parametric reference distribution of $\chi^2$ values. Comparison of the parametric and non-parametric distributions showed minimal differences

(Supplementary Fig. 14), confirming the robustness of our findings. The distribution of nonsynonymous mutation counts per gene was also examined to assess its skewness. We plotted the empirical distribution and observed that it was heavily skewed, with the majority of genes harbouring a low number of mutations, while a few genes contained disproportionately higher counts (Supplementary Fig. 15).

For each gene, the deviation from theoretical expectations was measured by their respective contribution to the $\chi^2$ statistic. The contribution of a gene $i$ was computed as percentage:

$$100\, x \frac{(n_i - mL_i)^2}{\chi^2_{obs}\, mL_i}$$

To identify genes that significantly deviate from the theoretical model, we implemented a sequential Jacknife procedure. This approach allowed us to iteratively exclude the gene with the highest contribution to the overall $\chi^2$ statistic and recompute the $\chi^2$ value after each exclusion. We continued this process until the $\chi^2$ statistic no longer showed a significant deviation from the theoretical distribution at a Bonferroni-corrected significance level. In total, this method identified n=60 genes exhibiting significantly higher nonsynonymous mutation counts than expected given their sequence length, under neutral evolution. The full list of these highly variable genes, along with their p-values, is provided in Supplementary Table 6. Code and data to generate the analysis is available at https://github.com/guillemmasfiol/GlobalPlague.

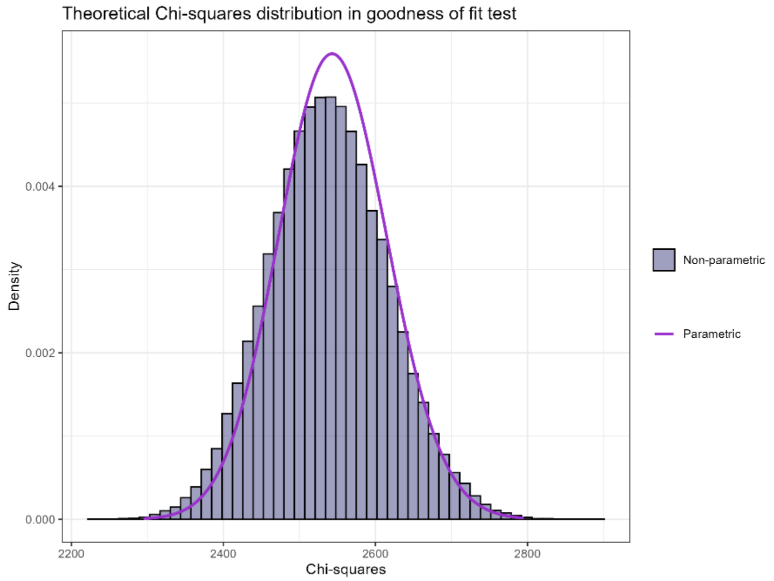

Supplementary Fig. 14 – Parametric and non-parametric (simulated) distributions for the $\chi^2$ values of the expected number of nonsynonymous mutations per gene.

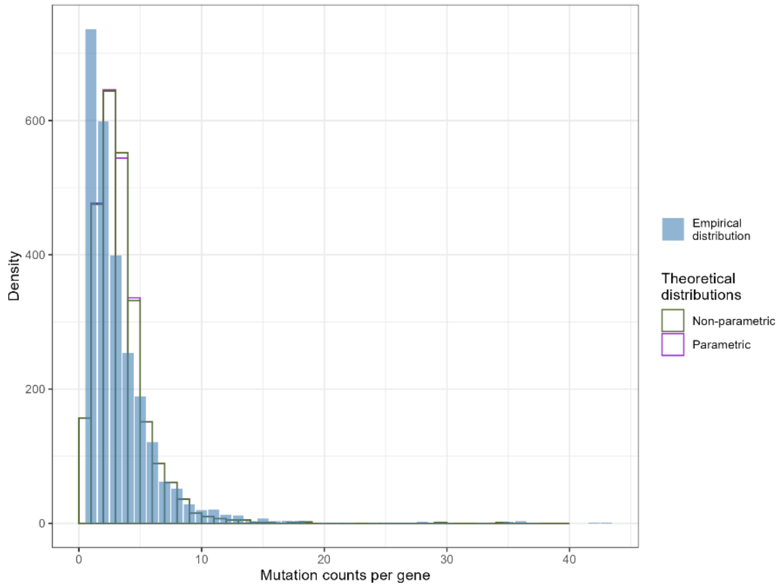

Supplementary Fig. 15 – Histograms showing the distributions of the observed (filled distribution) and theoretical (unfilled distributions) counts of nonsynonymous mutations per gene.

#### Supplementary Information 4

##### Supplementary Tables descriptions

All supplementary tables of this study are provided in spreadsheet format.

Supplementary Table 1 – Associated metadata of the 1,124 *Y. pestis* genomes sequenced in the study

Supplementary Table 2 – Table of the total 2,806 genomes included in the global phylogeny and associated metadata

Supplementary Table 3 – Summary table of diversity statistics (π and pairwise SNV distances) estimated for modern *Y. pestis* populations

Supplementary Table 4 – Comparative dataset of *Y. pestis* genomes included in the time-calibrated phylogeny inferred with BEAST 2.6.0

Supplementary Table 5 – Posterior distributions of divergence years for selected *Y. pestis* lineages inferred from the molecular dating analysis with BEAST 2.6.0

Supplementary Table 6 – Summary table of variant statistics and functional annotation (COG) of the 60 loci diversifying at significatively higher rates in *Y. pestis* genomes

Supplementary Table 7 – Summary of nonsynonymous and synonymous homoplastic mutation events per gene inferred using SNPPar 1.2

Supplementary Table 8 – Functional annotation of protein-coding genes in CO92 reference chromosome based on Clusters of Orthologous Groups (COG) using eggnog-mapper 2.1.2

Supplementary Table 9 – Predicted distances to DNA in mutated residues of RovA master virulence regulator and phylogenetic distribution of samples carrying mutated alleles

Supplementary Table 10 – Summary of plasmid sequences identified in *Y. pestis* genome assemblies using MOB-suite 3.0.1

Supplementary Table 11 – Identification of other mobile genetic elements (putative ICE and phage) across *Y. pestis* genome assemblies

Supplementary Table 12 – Total genes lost or inactivated through different evolutionary mechanisms (deletions, IS integration, nonsense mutations) across diverse modern *Y. pestis* genomes

Supplementary Table 13 – Genome annotations of the 20 Deletion Islands (DI) identified in modern and ancient *Y. pestis* chromosome and samples presenting each DI

Supplementary Table 14 – IS counts for the six IS types identified in the set of completed *Y. pestis* and *Y. pseudotuberculosis* genomes

Supplementary Table 15 – Total IS copy number per genome inferred from short-read data of modern *Y. pestis* genomes using ISMapper 2.0.2

Supplementary Table 16 – Total predicted IS integrations within each chromosomal gene identified in modern *Y. pestis* genomes

72. Tholozan, J. D. *Histoire de La Peste Bubonique En Perse*. (Masson, Paris, 1874).

73. Tholozan, J. D. *Histoire de La Peste Bubonique En Mésopotamie*. (G. Masson, Paris, 1874).

74. Tholozan, J. D. *Histoire de La Peste Bubonique*. (Masson, Paris).

75. Samedov, S. *Zdravokhraneniye v Iranskom Azerbaidzhane Nakanune i v Period Natsional'no-Osvoboditel'nogo i Demokraticheskogo Dvizheniya 1945-1946 Godov*. (Azerbaidzhanskoye Gosudarstvennoye Izdatel'svo, Baku, 1960).

76. Tholozan, J. D. *Les Trois Dernières Épidémies de Peste Du Caucase*. (Masson, Paris, 1879).

77. Hashemi Shahraki, A., Carniel, E. & Mostafavi, E. Plague in Iran: Its History and Current Status. *Epidemiol Health* e2016033 (2016).

78. Wickham, H. *Ggplot2: Elegant Graphics for Data Analysis*. (Springer-Verlag New York, 2016).
